# CytoPipeline & CytoPipelineGUI: A Bioconductor R package suite for building and visualizing automated pre-processing pipelines for flow cytometry data

**DOI:** 10.1101/2023.10.10.561699

**Authors:** Philippe Hauchamps, Babak Bayat, Simon Delandre, Mehdi Hamrouni, Marie Toussaint, Stéphane Temmerman, Dan Lin, Laurent Gatto

**Author notes:** Computational Biology and Bioinformatics, de Duve Institute, Université, Catholique de Louvain, Av., Hippocrate 75, 1200 Brussels, Belgium. **Data availability:** Raw flow cytometry data files, as well as manual gating information, are available on Zenodo (DOI:10.5281/zenodo.8425840). **Code availability:** All code needed to reproduce the results presented in the current article is available on the following GitHub repository: github.com/UCLouvain-CBIO/2023-CytoPipeline-code, of which a release has been archived on Zenodo (DOI:10.5281/zenodo.8425840). **Funding:** This work was funded by GlaxoSmithKline Biologicals S.A., under a cooperative research and development agreement between GlaxoSmithKline Biologicals S.A. and de Duve Institute (UCLouvain). **Competing interests:** B.B., S.D., M.H., M.T., S.T., and D.L. are employees of the GSK group of Companies. B.B., S.D., M.T., S.T., and D.L. report ownership of GSK shares. B.B., M.T., and S.T. are listed as inventors on patent(s) owned by the GSK group of companies. P.H. is a student at the de Duve Institute (UCLouvain) and participates in a post graduate studentship program at GSK.

## Abstract

**Background:** With the increase of the dimensionality in flow cytometry data over the past years, there is a growing need to replace or complement traditional manual analysis (i.e. iterative 2D gating) with automated data analysis pipelines. A crucial part of these pipelines consists of pre-processing and applying quality control filtering to the raw data, in order to use high quality events in the downstream analyses. This part can in turn be split into a number of elementary steps : signal compensation or unmixing, scale transformation, debris, doublets and dead cells removal, batch effect correction, etc. However, assembling and assessing the pre-processing part can be challenging for a number of reasons. First, each of the involved elementary steps can be implemented using various methods and R packages. Second, the order of the steps can have an impact on the downstream analysis results. Finally, each method typically comes with its specific, non standardized diagnostic and visualizations, making objective comparison difficult for the end user.

**Results:** Here, we present *CytoPipeline* and *CytoPipelineGUI*, two *R* packages to build, compare and assess pre-processing pipelines for flow cytometry data. To exemplify these new tools, we present the steps involved in designing a pre-processing pipeline on a real life dataset and demonstrate different visual assessment use cases. We also set up a benchmarking comparing two pre-processing pipelines differing by their quality control methods, and show how the package visualization utilities can provide crucial user insight into the obtained benchmark metrics.

**Conclusion:** *CytoPipeline* and *CytoPipelineGUI* are two Bioconductor *R* packages that help building, visualizing and assessing pre-processing pipelines for flow cytometry data. They increase productivity during pipeline development and testing, and complement benchmarking tools, by providing user intuitive insight into benchmarking results.

## Introduction

With recent advances in flow cytometry technologies, it has become possible to measure up to 50 markers simultaneously for the same single cells (McKinnon, 2018). As an immediate benefit, scientists now have access to richer flow cytometry experimental data. However, these advances also come at a cost, i.e. a more complex data analysis task. Indeed, traditional ‘manual gating’ data analysis procedures, which proceed by iterative hierarchical 2D representations of the data guided by biological knowledge, are unable to thoroughly extract the signal of interest from such high-dimensional data (Saeys, Van Gassen, and Lambrecht, 2016). There is therefore a need to complement such manual, expert-based approaches with *computational flow cytometry*, i.e. a set of computational algorithms and methods for automated, reproducible and data-driven flow cytometry data analysis (Saeys, Van Gassen, and Lambrecht, 2016).

Those computational flow cytometry approaches translate into so-called data analysis pipelines. Examples of such automated pipelines have been published in the recent litterature (e.g. Quintelier et al., 2021, Nowicka et al., 2017, Rybakowska et al., 2021, Ashhurst et al., 2022, Rybakowska et al., 2022). These consist of a series of data processing steps that are executed, one after the other, with the output of one step becoming the input of the next step. Schematically, for a typical flow cytometry data analysis, these numerous steps can usually be grouped into three big parts, coming after initial data sample acquisition:

- *data pre-processing and quality control*, which consists in both filtering undesirable and low quality events, and increasing the signal to noise ratio of the raw data, in order to feed the downstream steps with data of the highest possible quality;
- *population identification*, which aims at labelling the events with names of cell populations of interest;
- *downstream statistical analysis*, which can range from the simplest descriptive count/frequencies per population, to building complex prediction models for an outcome of interest, possibly for a high number of samples.

In what follows, we will mainly focus on the data pre-processing part, which can itself be split into several sub-tasks, or *steps* (Liechti et al., 2021): compensation, scale transformation, control (and possibly removal) of batch effects, control of signal stability in time (*QC in time*), filtering of undesirable events like debris, doublets and dead cells. All these steps are crucial to avoid that the subsequent analysis gets perturbed with erroneous signal (see e.g. Mazza et al., 2018 for compensation, Finak et al., 2010 for scale transformation, Emmaneel et al., 2022 for signal stability in time, and den Braanker, Bongenaar, and Lubberts, 2021 for other steps).

However, building good, automated pre-processing pipelines, suitable for the particular type of biological samples and biological question, can be challenging for a number of reasons. First, for each elementary step, there might exist a number of different computational methods, each of those having numerous parameters available to the user. For example, looking only into methods available on the Bioconductor project (Huber et al., 2015) for controlling the signal stability (*QC in time*), one finds at least four different methods available: *flowAI* (Monaco et al., 2016), *flowClean* (Fletez-Brant et al., 2016), *PeacoQC* (Emmaneel et al., 2022), *flowCut* (Meskas et al., 2023), and each of these methods comes with 7 to 11 different parameters. Second, the order of steps is not always set in stone, and applying different orders can lead to different outcomes, an effect coined steps interaction. These two facts lead to what we refer to as the *combinatorial problem of designing pipelines*, which means that, as the number of necessary elementary steps increases, the number of possible pipeline designs grows exponentially. As a consequence, for the user, it becomes time consuming to build and assess even only a few of the possible step combinations, let alone testing a representative sample of them in a systematic manner.

On top of that combinatorial problem, the user is also faced with a lack of generic, standardized and user-friendly tools to evaluate and compare data pre-processing pipelines. On the one hand, each single pre-processing step method might come with its own approach for diagnostic and visualization (e.g. ad hoc plots, html or pdf reports), which allows the user neither to standardize the comparison process, nor to easily investigate the links and interactions between the different steps. On the other hand, there are a number of benchmarking studies comparing computational methods for flow cytometry data (Liu et al., 2019, Weber and Robinson, 2016, Cheung et al., 2022, Aghaeepour et al., 2016), but they tend to focus on only one part of the pipeline, which is usually the downstream analysis. Finally, there also exist some generic tools and frameworks to systematize the benchmarking process, including the interactions between different steps, such as for example *pipeComp* (Germain, Sonrel, and Robinson, 2020) and *CellBench* (Su et al., 2020), and as well as some attempts to formally model the pipeline optimization problem in mathematical terms (Selega and Campbell, 2022). However, what is lacking for the end user is the ability to intuitively interpret the results of such benchmarkings. In other words, could one translate that a pipeline *A* outperforms a pipeline *B*, with respect to a specific performance metric, in terms of the obtained data characteristics, or number of filtered events. Therefore, there is still a need for flow cytometry practitioner-focused standardized tools for visual comparison of pre-processing pipelines.

Here, we present *CytoPipeline* and *CytoPipelineGUI*, two *R* packages aimed at facilitating the design and visual comparison of pre-processing pipelines for flow cytometry data. We describe the concepts underlying the software, provide some illustrative examples and demonstrate the use of the accompanying visualization utilities. We show that these new tools can help increasing the productivity during pipeline development and testing, and that they can complement benchmarking tools and studies, by providing the user with intuitive insight into benchmarking results. The *CytoPipeline* and *CytoPipelineGUI* packages are available on Bioconductor (Huber et al., 2015), as of version 3.17 and 3.18, respectively.

## Methods

### Implementation

In what follows, we assume that we have a dataset, provided as a set of files in Flow Cytometry Standard (*fcs*) format (Spidlen et al., 2010), on which we would like to apply a data pre-processing pipeline.

The *CytoPipeline* suite is composed of two main *R* packages, *CytoPipeline* and *CytoPipelineGUI*. While *CytoPipeline* is the main package, providing support for pipeline definition, running, monitoring and basic ploting functions, *CytoPipelineGUI* provides two interactive GUI applications enabling users to interactively explore and visualize the pipeline results. The *CytoPipeline* framework is based on two main concepts, namely *CytoPipeline* and *CytoProcessingStep*. A *CytoPipeline* object centralizes the pipeline definition, and specifies the run order of the different pipeline steps. These steps materialize as *CytoProcessingStep* objects, which store pipeline step names and the corresponding *R* functions that will be called at execution time. These functions are either provided within the *CytoPipeline* package itself, exported from third party packages, or coded by the user themself. Together with the function name to be called, a *CytoProcessingStep* object also contains the list of parameters that are used as arguments to the function.

When creating a *CytoPipeline* object, the user can provide the description of the pipeline as a text file in *json* format (Pezoa et al., 2016). Figure 1 shows the typical structure of such a *json* file. Note that, in practice, two different sets of processing steps, or pipelines, are described:

- A *scaleTransformProcessingSteps* pipeline, which describes the set of successive steps needed to generate the scale transformations that will be applied to the different channels of each of the *fcs* files that are included in the dataset. The *CytoPipeline* engine will run this pipeline first, and only once, prior to running the pre-processing on each *fcs* file.
- A *flowFramesPreProcessingSteps* pipeline, which describes the set of pre-processing steps that will be applied on each of the different *fcs* file independently.

**Figure 1.**
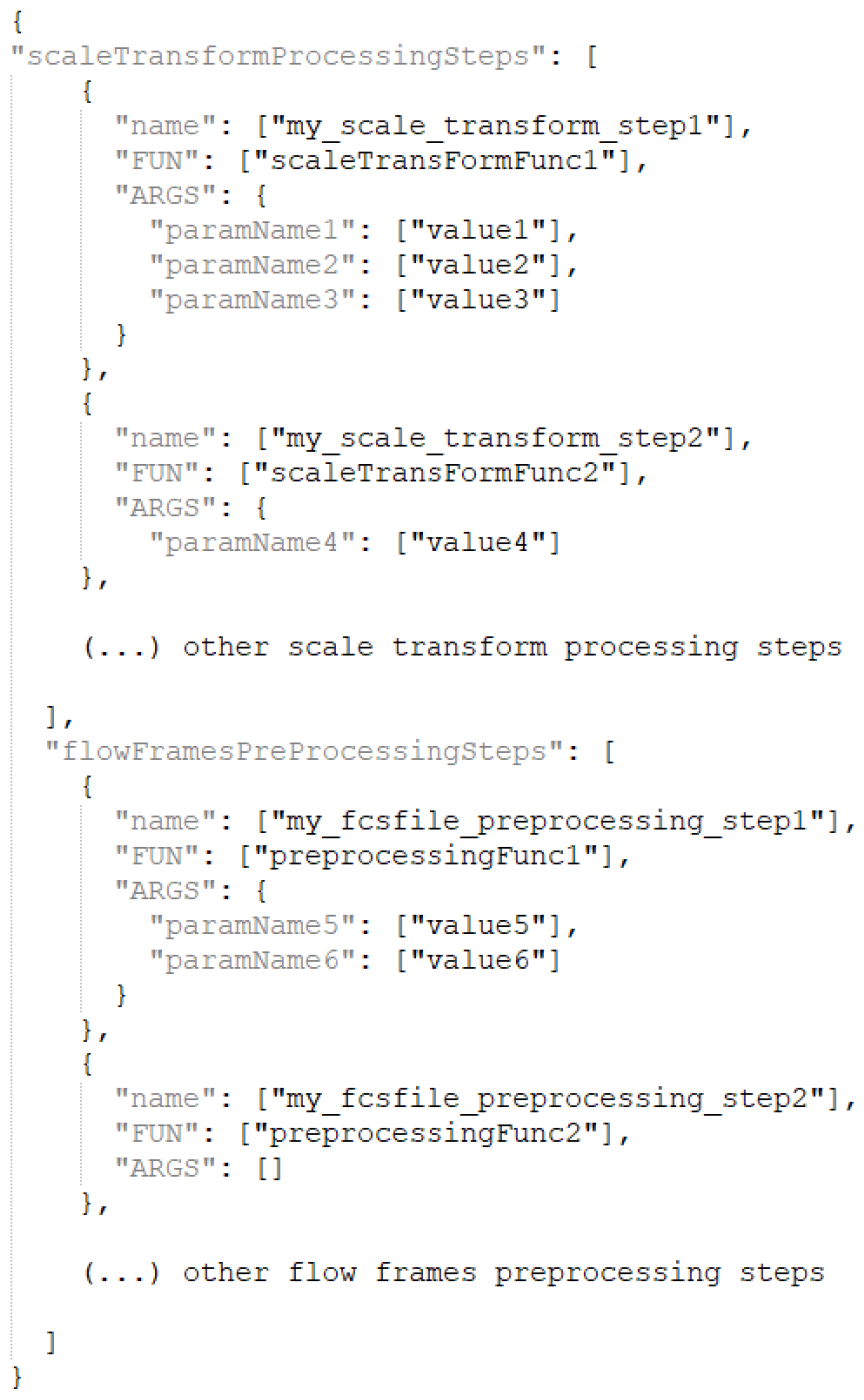
Structure of the user provided json file that describes a *CytoPipeline* object. The first pipeline (i.e. *“scaleTransformProcessingSteps”*) specifies how the preliminary calculation of the scale transformations is performed. Here only its first two steps are described. The first step, named *“my_scale_transform_step1”*, consists in calling the *“scaleTransFormFunc1”* function, with 3 parameters (*“paramName1”, “paramName2”* and *“paramName3”* provided as arguments, taking the specified *“value1”, “value2”* and *“value3”* value respectively. The second step, named *“my_scale_transform_step2”*, calls the *“scaleTransFormFunc2”*, with only one single parameter, i.e. *“paramName4”* taking *“value4”* as value. The second pipeline (i.e. *“flowFramesPreProcessingSteps”*) specifies the set of pre-processing steps performed on each data file independently. Here again, only the first two steps are described. The first one, named *“my_fcsfile_preprocessing_step1”* calls the *“preprocessingFunc1”* function with two parameters, while the second one, named *“my_fcsfile_preprocessing_step2”* calls the *“preprocessingFunc2”* function with no parameter (apart from the output of the previous step which is always used as an implicit additional argument).

Steps in both pipelines are described in the exact same way, i.e. by providing a user-chosen name for the step, the corresponding function that needs to be called by the engine upon running, and the set of arguments (i.e. the list of parameter names and corresponding values) that need to be provided to the function. Note that, on top of these explicitely defined arguments, the running engine will also take the output of each step as an implicit additional argument to the function called by the subsequent step.

The standard process for using *CytoPipeline* to build, run and inspect pre-processing pipelines is the following:

- define the pipeline by specifying the different steps using a descriptive text file, in *json* format^1^;
- run the pipeline, possibly for several data files in parallel, which involves writing and executing a short *R* script (see following sections);
- monitor the execution process thanks to a *CytoPipeline* provided workflow visualization utility;
- visualize and compare the results at different stages, using the *CytoPipelineGUI* interactive GUI applications.

In terms of technical infrastructure, the *CytoPipeline* package suite makes itself internal use of several technical *R* packages:

- *BiocParallel* (Morgan et al., 2023) enabling parallel pre-processing of *fcs* sample files;
- *BiocFileCache* (Shepherd and Morgan, 2023) enabling storage (i.e. *caching*) of all intermediary results for further inspection;
- *shiny* (Chang et al., 2023) for interactive visualizations.

### Illustrative dataset

In order to demonstrate *CytoPipeline* functionalities, we make use of an illustrative dataset, the *HBV chronic mouse* dataset. This dataset was collected during a preclinical study aimed at assessing the effect of different therapeutic vaccine regimens on the immune response of Hepatitis B Virus transduced mice.

In this study, 56 male and female HLA.A2/DRB1 transgenic mice (transgenic for the human HLA-A2 and HLA-DRB1 molecules) were used. HLA.A2/DRB1 mice from groups 1, 2 and 4 were transduced at day 0 with adeno-associated virus serotype 2/8 (AAV2/8-HBV) vector carrying a replication-competent HBV DNA genome and randomized before immunization with 4 doses of vaccine candidate, based on level of HBs circulating antigen detected in the sera at day 21, age and gender proportions. Mice from group 3 were not transduced with AAV2/8-HBV viral vector and were immunized with four doses of vaccine candidate and finally, mice from group 5 were not transduced and received four doses of NaCl solution. Upon sacrifice, livers were collected, perfused with Phosphate Buffered Saline (PBS) to remove blood cells and after enzymatic treatment, lymphocytes were isolated, stained with different monoclonal antibodies. The stained cells were acquired by flow cytometry using a BD Symphony A5 flow cytometer - the same instrument for all biological samples - and analyzed using the FlowJo v10.8 Software (BD Life Sciences).

Animal husbandry and experimental procedures were ethically reviewed and carried out in accordance with European Directive 2010/63/EU and the GlaxoSmithKline Biologicals’ policy on the care, welfare and treatment of animals, in GSK animal facilities located in Rixensart, Belgium (AAALAC accredited). The ethical protocol of the GSK in vivo study was approved by the local GSK ethical committee.

This experiment resulted in the acquisition of 55 different *fcs* raw data file - one sample could not be acquired - with a flow cytometry panel of 12 different channels. The *HBV chronic mouse* dataset is available on Zenodo (DOI:10.5281/zenodo.8425840).

### Applied pre-processing pipelines

#### Pipeline set-up

For the purpose of illustrating *CytoPipeline* functionalities, the 55 raw data files of the *HBV chronic mouse* dataset were used as input of two different pre-processing pipelines. Each pipeline was composed of the following steps:

- Reading of the raw *fcs* sample files, using the *flowCore* package (Ellis et al., 2023).
- Margin events removal, which consists in identifying and removing the outliers using the *PeacoQC* package (Emmaneel et al., 2022). In short, manual boundaries per channel, corresponding to the instrument detection limits, are applied, and all events falling outside these boundaries are removed.
- Signal compensation, which consists in applying an existing compensation matrix. This matrix was generated by the flow cytometer at data acquisition time, and subsequently manually adjusted by the expert scientist.
- *QC in time*, which consists in eliminating parts of the signal that are not stable in time, using one of the corresponding QC algorithms (see below).
- Doublet removal, which consists in keeping the events that have a similar area vs. height ratio of the FSC channel signal pulse, and eliminating the doublets, which have a significantly higher ratio. This was performed using an ad hoc implementation in the *CytoPipeline* package.
- Debris removal, which consists in clustering the events in the (FSC-A, SSC-A) 2D representation, targetting a number of clusters provided by the user. After the clusters are obtained, the cluster of which the centroid lies nearest to the origin, i.e. with the smallest FSC-A (size) and smallest SSC-A (content, granularity), is considered as containing debris and removed. This was done using the *flowClust* package (Lo et al., 2009).
- Scale transformation, which consists in automatically estimating the parameters of a *logicle* transformation (Parks, Roederer, and Moore, 2006), using the *flowCore* package (Ellis et al., 2023). The obtained scale transformations were applied on all 55 sample files, and the parameters were estimated on an aggregation of a subset of 4 randomly chosen sample *fcs* files, after margin events removal and signal compensation.
- Dead cells removal, which consists in automatically setting a threshold between live cells and dead cells in the corresponding fluorescent ‘Live & Dead’ channel dimension, using the *flowDensity* package. The events having a ‘Live & Dead’ intensity above the found threshold are eliminated as dead cells.

However, the two pre-processing pipelines essentially differed by the method used for the *QC in time* step, as one used the *PeacoQC* package (Emmaneel et al., 2022), while the other used the *flowAI* package (Monaco et al., 2016). In addition, the step order was also different, as the *PeacoQC* method is based on a peak detection algorithm which needs to run on compensated, scaled transformed data (Emmaneel et al., 2022), while the *flowAI* method is advised to be applied on raw data (Monaco et al., 2016). Figure 2 outlines the different steps applied in the pre-processing of each *fcs* files, for both the *PeacoQC*-based pipeline, and the *flowAI*-based pipeline.

**Figure 2.**
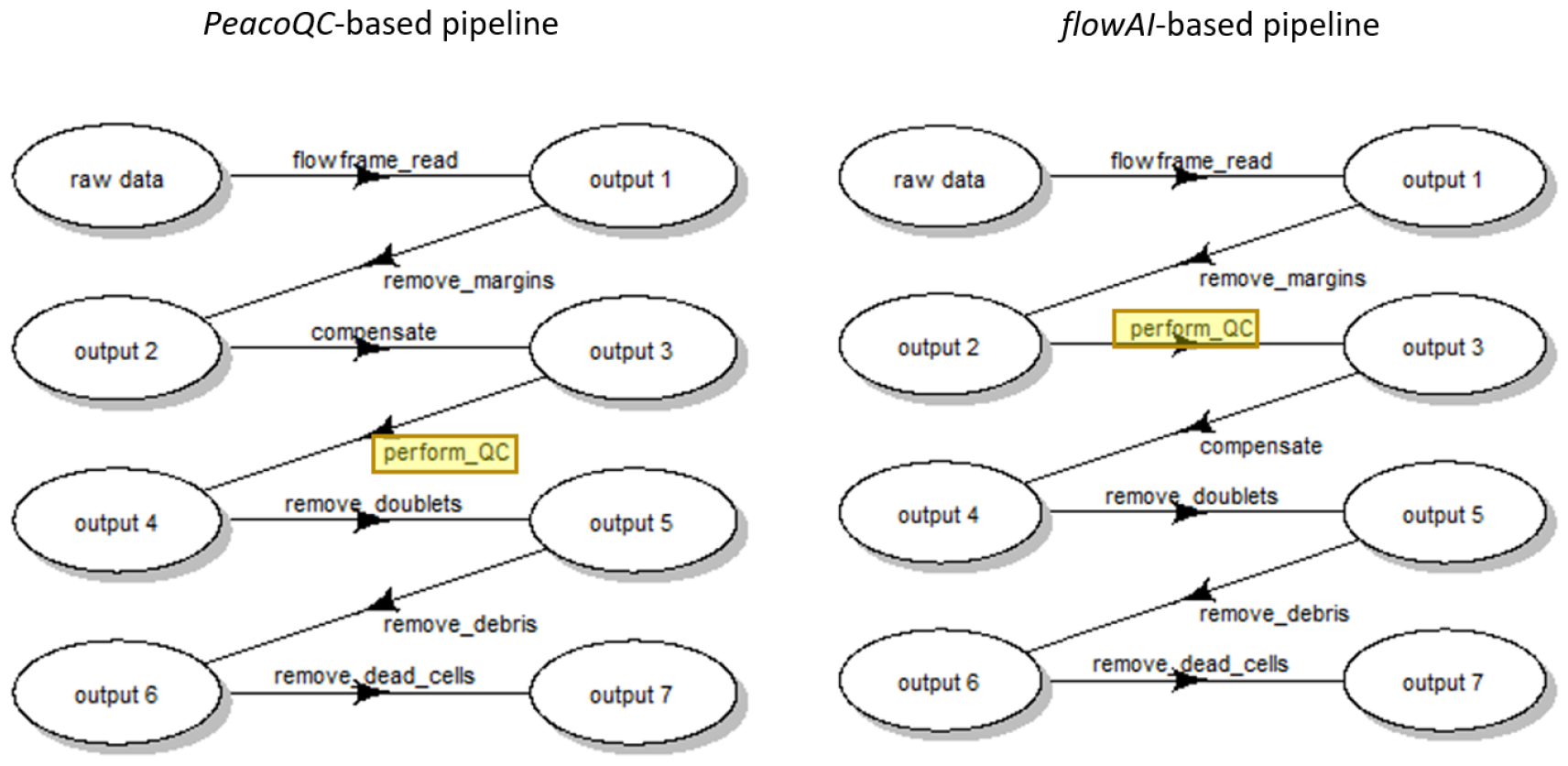
Workflow of the subsequent steps applied in the pre-processing of each fcs file, for both pipelines. Note that these plots have been generated using the *CytoPipeline* package.

More detailed information on packages, versions and methods underlying each step (Tables S1, S2 & S3), as well as the *json* configuration files defining respectively the *PeacoQC*-based and *flowAI*-based pipelines are available in the supplementary materials.

#### Running the pipelines and visualizing the results

In order to create the *CytoPipeline* objects representing the pipelines, run them and visualizing the results - including monitoring of the pipeline execution - a short *R* script needs to be written and executed. An example of such *R* script is provided in Figure S1. Note that, as a result of the centralization of the pipeline definition, the code is very simple and concise, as for example, creating and running the pipeline boils down to essentially two *R* statements. Also, note that it is the same *R* code that triggers the execution of both pre-processing pipelines described in the previous section (except for the selection of the appropriate input *json* file and the choice of the experiment name under which to store the results). The distinctive part of the pipeline is located in the input *json* file, which describes the pipelines steps and their execution order.

### Example benchmarking

Aiming at illustrating the use of *CytoPipeline* to provide insights into benchmarking results, we designed a benchmarking, which consisted in comparing the outcome of the two competing *PeacoQC*-based and *flowAI*-based pipelines described in the previous section, using the *HBV chronic mouse* dataset, to a ground truth. The latter was obtained by submitting the 55 raw data *fcs* files to an expert scientist, who manually pre-processed the files, gated the events using FlowJo. The obtained FlowJo workspace file was subsequently automatically processed using the *CytoML* package (Finak, Jiang, and Gottardo, 2018) version 2.12.0, and incorporated into a dedicated *CytoPipeline* ground truth pipeline for comparison with the two automated pipelines.

Regarding the benchmark evaluation metrics, for each single *fcs* file, the final output of each pipeline was compared to the ground truth, in terms of number of events, and the following metrics were calculated: sensitivity, specificity, precision and recall, which are defined as follows:

let

- *G* (resp. *B*) be the set of events that are considered as **G**ood (resp. **B**ad) in the manual gating i.e. in the ground truth;
- *FG* (resp *FB*) be the set of events that are flagged as good (resp. flagged as bad) by the considered automated pipeline.

We can additionally define the following sets of events:

- *FG*, correct *= FG ∩ G*, the set of events that are correctly flagged as good;
- *FB*, correct *= FB ∩ B*, the set of events that are correctly flagged as bad.

The chosen evaluation metrics are then defined as:

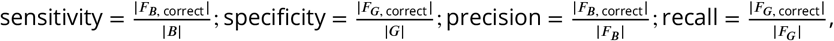

where |*A*| stands for the number of elements in the set A.

The benchmark was set up and performed using the *pipeComp* package (Germain, Sonrel, and Robinson, 2020), version 1.10.0. Indeed, *pipeComp* is a convenient tool to efficiently automate multiple alternative pipelines to be compared in the benchmark, as well as to automate the calculation of the evaluation metrics for each dataset used as benchmark input.

## Results

### Visual assessment and comparison of pipeline outputs

We used *CytoPipeline* to define both *PeacoQC*-based and *flowAI*-based pre-processing pipelines, as described in the Methods section, on the *HBV chronic mouse* dataset. We obtained results in the form of sets of data matrices (or *flowFrames*) after each step for each pre-processing pipeline. In the following paragraphs, we present some *CytoPipeline* visual assessment plots, according to 6 different use cases (Table 1). Use case #1 consists in visualizing a run and monitoring the status of the different steps. Use cases #2 to #5 consist in either looking at ‘what happened’ within a single pipeline for a single biological sample in isolation (use case #2), or comparing two different situations (flow frames) involving different pipelines (use cases #3 and #4), or involving different biological samples within the same pipeline (use case #5). Finally, use case #6 consists in assessing, and possibly modifying, the scale transformations obtained during a pipeline execution.

**Table 1.** Use cases of visual assessment and comparison of pipeline outputs. When the use case involves comparing two flow frames obtained from different steps and/or different pipelines (i.e. use cases #2 to #4), or different samples (i.e. use case #5) the 3 columns ‘sample’, ‘pipeline’ and ‘output’ designate the initial flow frame (referring to Figure 2), while the 3 columns ‘compared sample’, ‘compared pipeline’ and ‘compared output’ designate the flow frame that is compared to the initial flow frame.

| Use case | Description | Sample | Pipeline | Output (cf. Figure 2) | Compared sample | Compared pipeline | Compared output (cf. Figure 2) |
| --- | --- | --- | --- | --- | --- | --- | --- |
| #1 | Monitoring a run | D91_G01 | PeacoQC | All | Not applicable | Not applicable | Not applicable |
| #2 | Visualizing the effect of a single pipeline step | D91_G01 | PeacoQC | output 2 (before ‘compensate’) | D91_G01 | PeacoQC | output 3 (after ‘compensate’) |
| #3 | Comparing the outcome of a pipeline step with different parameter values | D91_G01 | PeacoQC | output 6 (after ‘remove_debris’, run with 3 clusters) | D91_G01 | PeacoQC | output 6 (after ‘remove_debris’, run with 2 clusters) |
| #4 | Comparing two different methods for one or several step(s) | D91_A01 | PeacoQC | output 7 (after ‘remove_dead_cells’) | D91_A01 | flowAI | output 7 (after ‘remove_dead_cells’) |
| #5 | Comparing two different biological samples | D91_A01 | PeacoQC | output 7 (after ‘remove_dead_cells’) | D93_B05 | PeacoQC | output 7 (after ‘remove_dead_cells’) |
| #6 | Visualization and update of generate scale transformations | D91_G01 | PeacoQC | Not applicable | Not applicable | Not applicable | Not applicable |

#### Use case #1: Monitoring a run

As all the intermediate results produced during pipeline execution are saved (see Methods/Implementation section), it is possible to generate a summary workflow view, consecutive to a run. Figure 3 shows an example of such a display, obtained after running the *flowAI*-based pipeline described above, where there was a spelling error in one of the parameter names of the *“remove_debris”* step. On top of showing the sequence of steps, a colour code is used to highlight which of the steps have run to completion, and which of the steps need to be re-run. Here, for the selected sample, the pipeline ran correctly until the *“remove_doublets”* step (green nodes), but did not produce any output for the subsequent steps (orange nodes), which is due to the spelling error in the definition of the *“remove_debris”* step. Based on this summarized visual information, the user can now dig into the flagged problematic step, and/or track the particular characteristics of the sample which generated the error. More details on the colour code used in this plot can be found in Figure S2.

**Figure 3.**
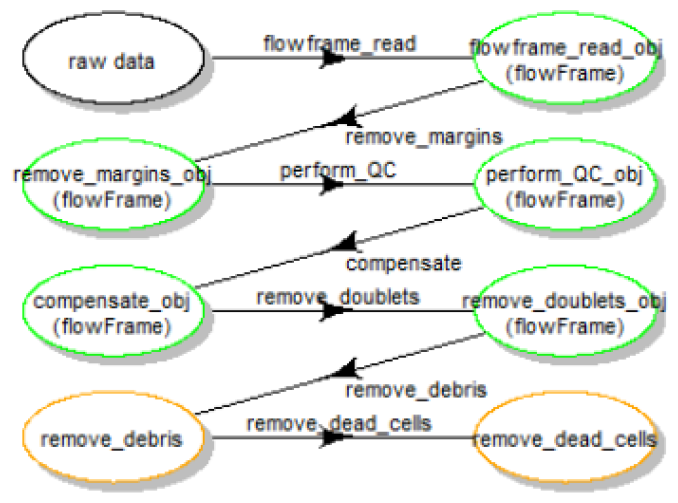
Use case #1: summary workflow view of the run - green nodes correspond to steps that ran to completion for the selected sample file, orange nodes correspond to steps that have not generated an output yet.

#### Use case #2: Visualizing the effect of a single pipeline step

In Figure 4, the user is visually assessing two consecutive states of the *flowFrame* of sample *D91_G01*, within the same run of the *PeacoQC*-based pipeline. To evaluate the effect of the compensation step, the *“before compensation”* (output 2, cf. Figure 2) and the *“after compensation”* (output 3) states of the pipeline are visually compared. Note that this visualization can be done according to any pair of selected channels/markers (2D distribution representation), or according to a 1D marginal distribution representation for any selected channel/marker. Here, the (CD8, CD38) 2D view shows, on the left, that the fluorescence of the dye BB700 (CD38) spills into the CD8 channel. On the right, application of a pre-computed compensation matrix (see Methods section) has rectified the distribution of the two markers, revealing different ranges of CD38 (an activation marker) between the CD8+ and CD8-populations. Note that a corresponding screenshot of the interactive GUI application, implemented in the *CytoPipelineGUI* package, can be found in Figure S3.

**Figure 4.**
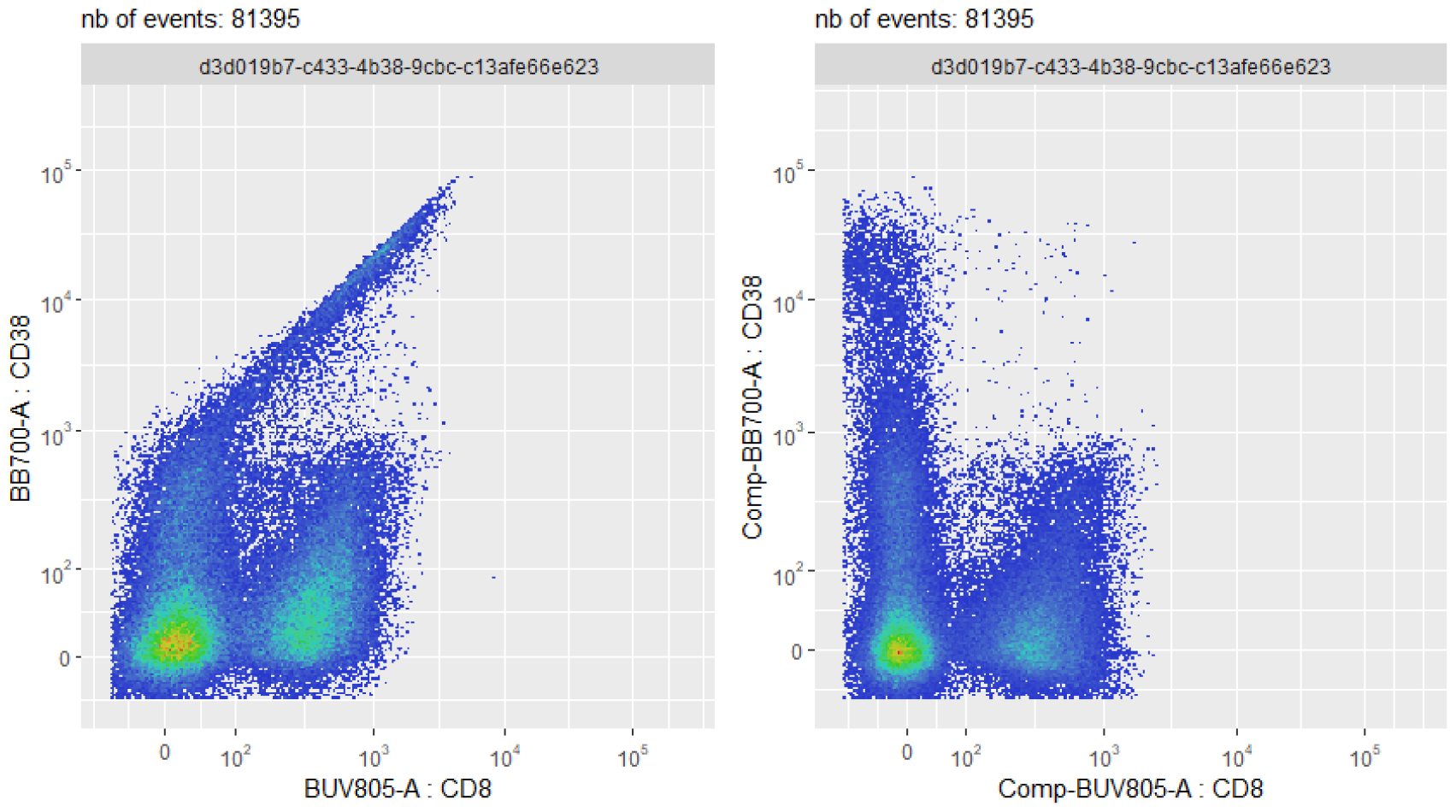
Use case #2: effect of a single pipeline step - here, the compensation step of the *PeacoQC*-based pipeline for sample *D91_G01*. On the left, spillover of the BB700 (CD38) dye fluorescence into the CD8 channel creates a visual artefact, with events wrongly flagged as double positive CD8+ CD38+. On the right, compensation has rectified the bivariate distribution of the two markers.

#### Use case #3: Comparing the outcome of a pipeline step with different parameter values

This use case involves running the same pipeline, with the same steps but with amended values for one or several steps, in order to investigate which parameter combination performs better. An illustrating example is shown in Figure 5, where the outcome of the debris removal step is compared when applying two different user input number of clusters (three on the left plot, vs. two on the middle plot). On the right plot, events coloured in red are the ones that are eliminated when applying the debris removal step when the number of clusters is two, but not eliminated when the number of clusters is three. Let us recall that the debris elimination step consists in clustering the events in a fixed number of clusters, followed by the elimination of the cluster nearest to the origin - see Methods section. Here, specifically, the user can conclude that the debris removal algorithm (based on *flowClust* package) does a better job selecting the target events when the appropriate number of target clusters is used, i.e. two clusters, as on the middle plot. This is because the cell population of interest, here a population of lymphocytes extracted from mice liver tissues, naturally groups into one single cluster in the (FSC-A, SSC-A) 2D representation. As a consequence, in this case, two is the optimal number of clusters (one cluster of debris, one cluster of lymphocytes). Figure S4 illustrates the removal of events during the debris removal step, for the 2 clusters and the 3 clusters cases.

**Figure 5.**
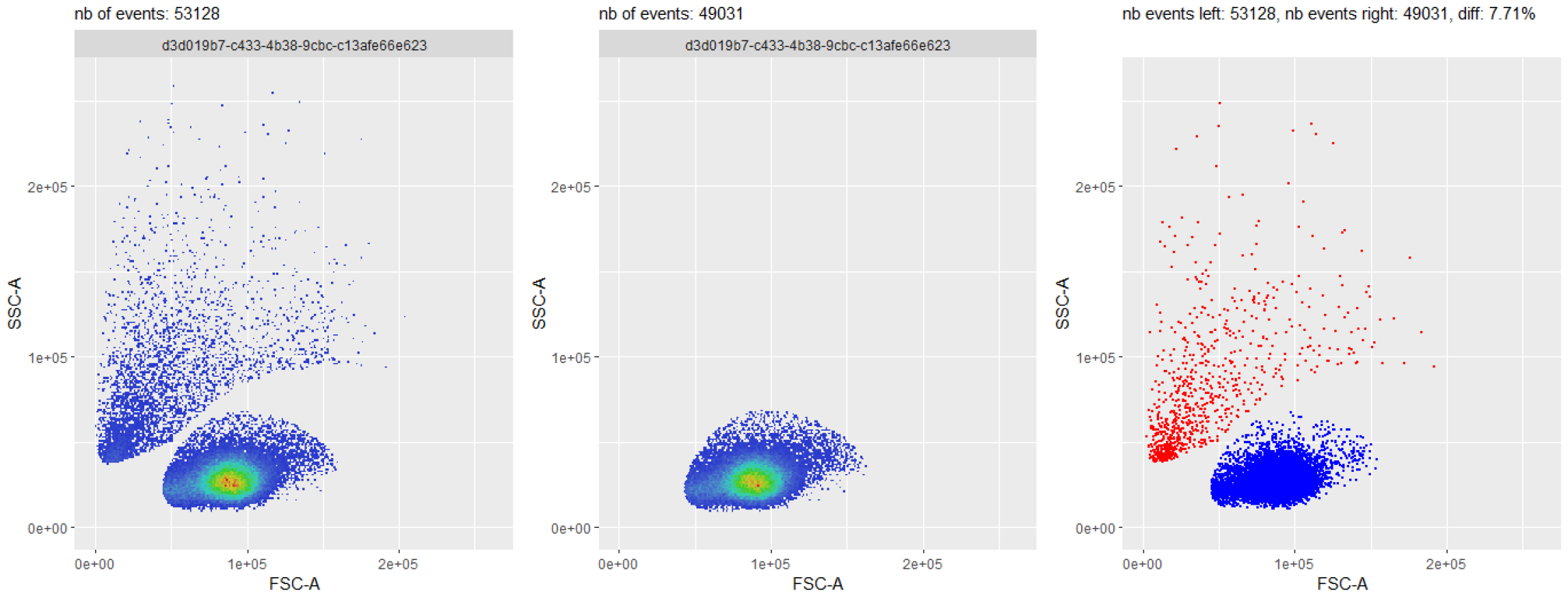
Use case #3: comparison of two different parameter settings for the debris removal step, on sample *D91_G01*. The setting with two clusters (in the middle) better eliminates undesirable events than the setting with three clusters (on the left). On the right, an explicit comparison between the two flowFrames is performed. Red dots correspond to events that are present on the left hand side plot, but not present on the middle plot, while blue dots correspond to events that are present on both plots.

#### Use case #4: Comparing two different methods for one or several steps

This use case is a generalization of the preceeding one, where the user wants to compare the performance of two different methods for one or several steps of the pre-processing pipeline. For instance, Figure 6 provides a comparison between the *PeacoQC*-based and the *flowAI*-based pipelines, applied on a particular biological sample of the *HBV chronic mouse* dataset. This comparison, obtained by plotting one specific channel (here the *FSC-A*) as a function of time, reveals that *flowAI* removes time chunks more aggressively than *PeacoQC*, for the current sample. Note that this comparison can also be done for any 2D combination of makers (not shown here).

**Figure 6.**
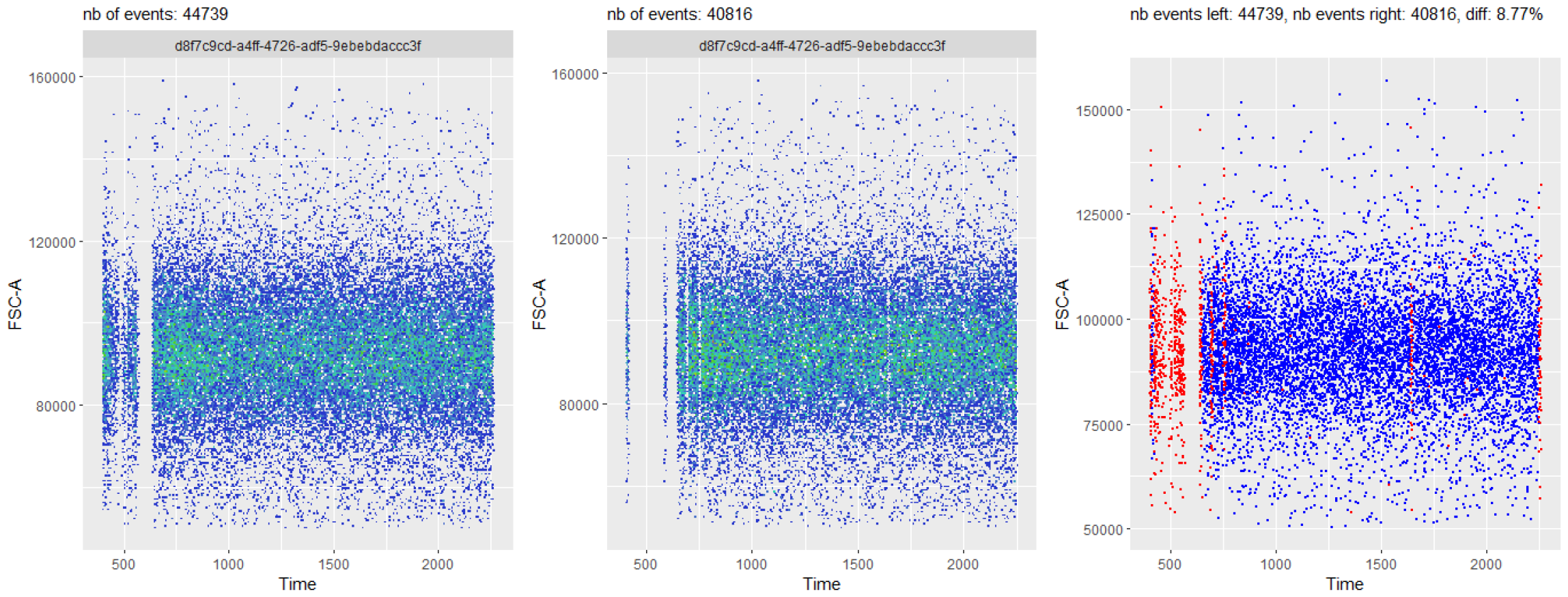
Use case #4: comparison of the final state results, for sample *D91_A01*, between the *PeacoQC*-based pipeline and the *flowAI*-based pipeline - here using a (*FSC-A* vs. *Time*) plot. On the left, the end state of the *PeacoQC*-based pipeline is shown, while the end state of the *flowAI*-based pipeline is shown on the middle plot. On the right, an explicit comparison between the two flowFrames is performed. Red dots correspond to events that are present on the left hand side plot, but not present on the middle plot, while blue dots correspond to events that are present on both plots. This figure reveals that, for this particular sample, *flowAI* tends to remove time chunks more aggressively than *PeacoQC*.

#### Use case #5: Comparing two different biological samples

It is also possible to compare two different biological samples of the same dataset, at any specific step of any pipeline. This allows e.g. to check that the methods used for the various pre-processing steps perform consistently across the whole dataset. One example is shown in Figure 7, where two different samples are displayed in a 2D plot with the *FSC-A* and *Live&Dead* channels. In this case, the two samples show very similar bivariate distributions. Based on this 2D representation, one could conclude that the pre-processing pipeline has correctly selected the target cell population in both cases. This would however need careful confirmation based on other 2D combinations of markers, e.g. *FSC-H* vs. *FSC-A* (for doublets elimination), and *FSC-A* vs. *SSC-A* (for debris elimination).

**Figure 7.**
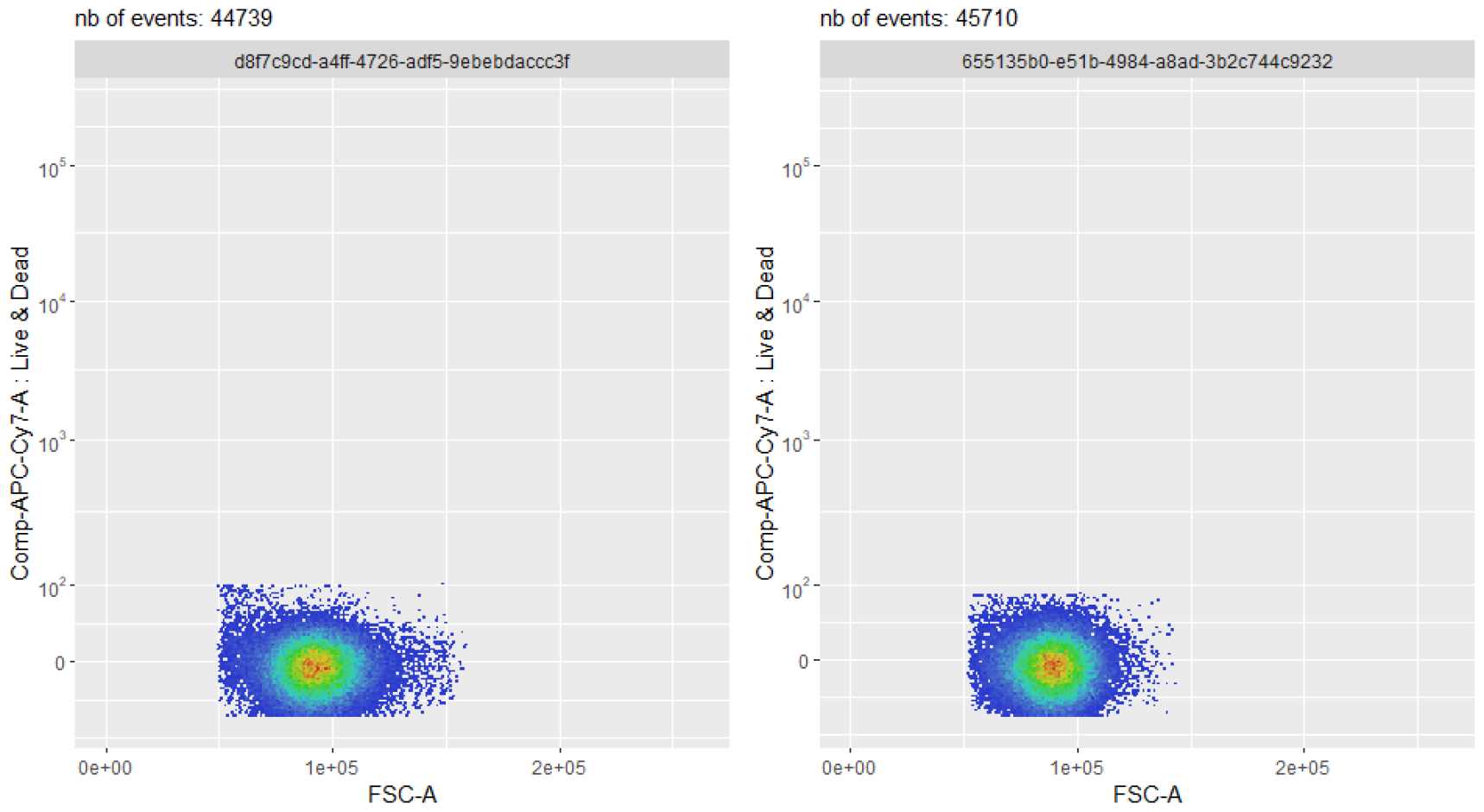
Use case #5: comparison of the final state results between two different biological samples (on the left: sample *D91_A01* and on the right: sample *D93_B05*), within the same *PeacoQC*-based pipeline. For the particular channels chosen (*Live&Dead* vs. *FSC-A*), the two samples show very similar bivariate distributions. Based on this 2D representation, one could conclude that the pre-processing pipeline has correctly selected the target cell population in both cases.

#### Use case #6: Visualization and update of generated scale transformations

Besides the *flowFrame* comparison tool, *CytoPipelineGUI* also provides a second interactive GUI application, which is aimed at inspecting the scale transformations obtained from the corresponding *scaleTransformProcessingSteps* pipeline (see Methods/Implementation section). If the shape of the distribution after transformation needs adjustment (for example for better separation of negative and positive populations for a specific marker), the user can manually adapt the scale transformation parameters, interactively assess the impact of their modifications, and apply these modifications to the scale transformations for further use in the pre-processing pipelines (Figure 8). These manual adjustments can be very useful, for example when the automatic transformation parameter adjustment algorithm has not worked satisfactorily. Figure 9 shows an example where the *logicle* transformation (Parks, Roederer, and Moore, 2006) applied on marker CD38 (left) shows spurious density oscillations in the negative domain. Manually adjusting the *positive decimals* parameter of the *logicle* transformation leads to a better looking density plot, where one can more easily distinguish CD38-, CD38+ and CD38++ populations.

**Figure 8.**
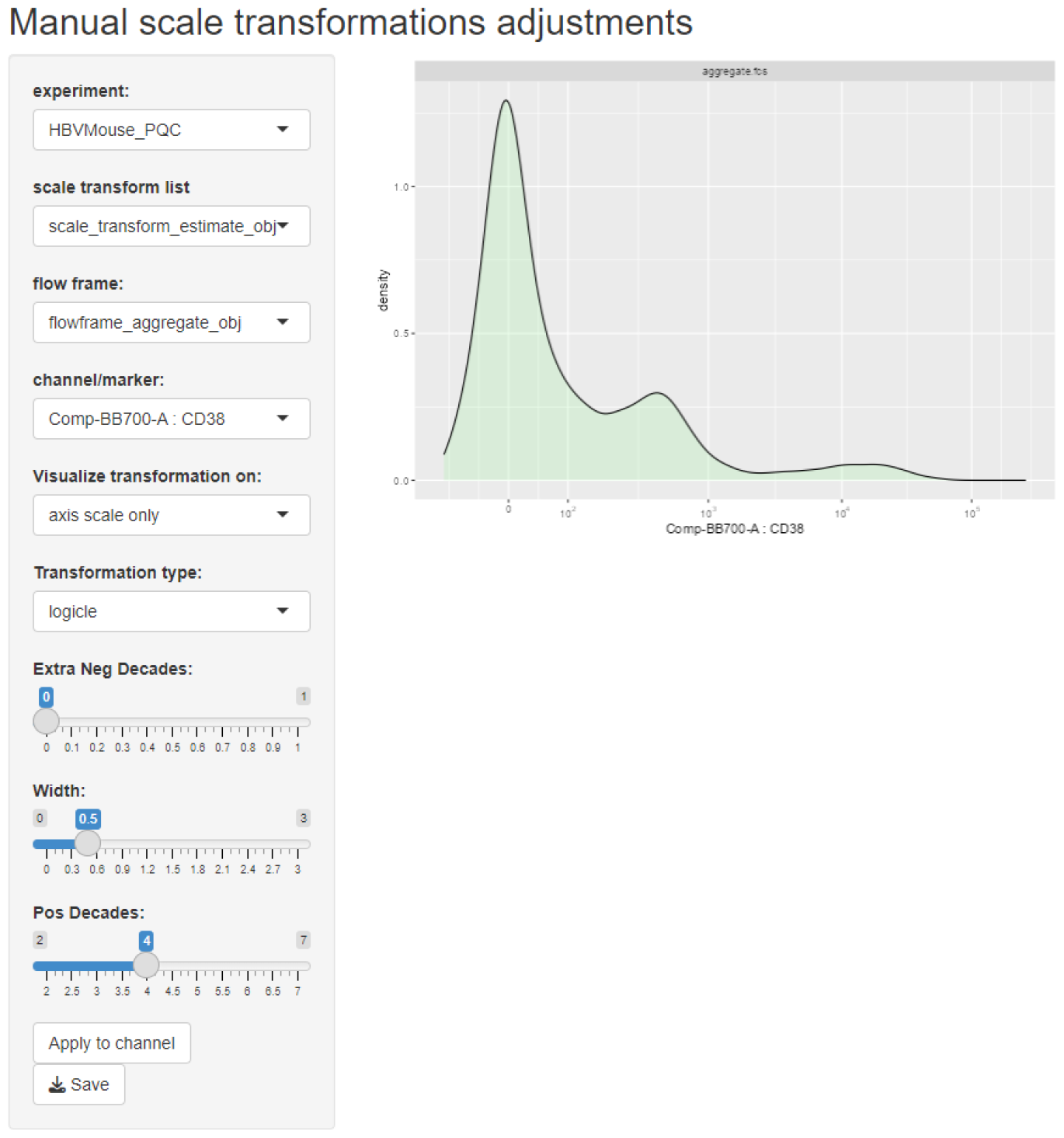
Use case #6: screenshot of the *CytoPipelineGUI* interactive GUI application enabling the inspection, manual adjustment and save of pipeline generated scale transformations. Here the user is visualizing the transformation applied on marker CD38, for sample *D91_G01*

**Figure 9.**
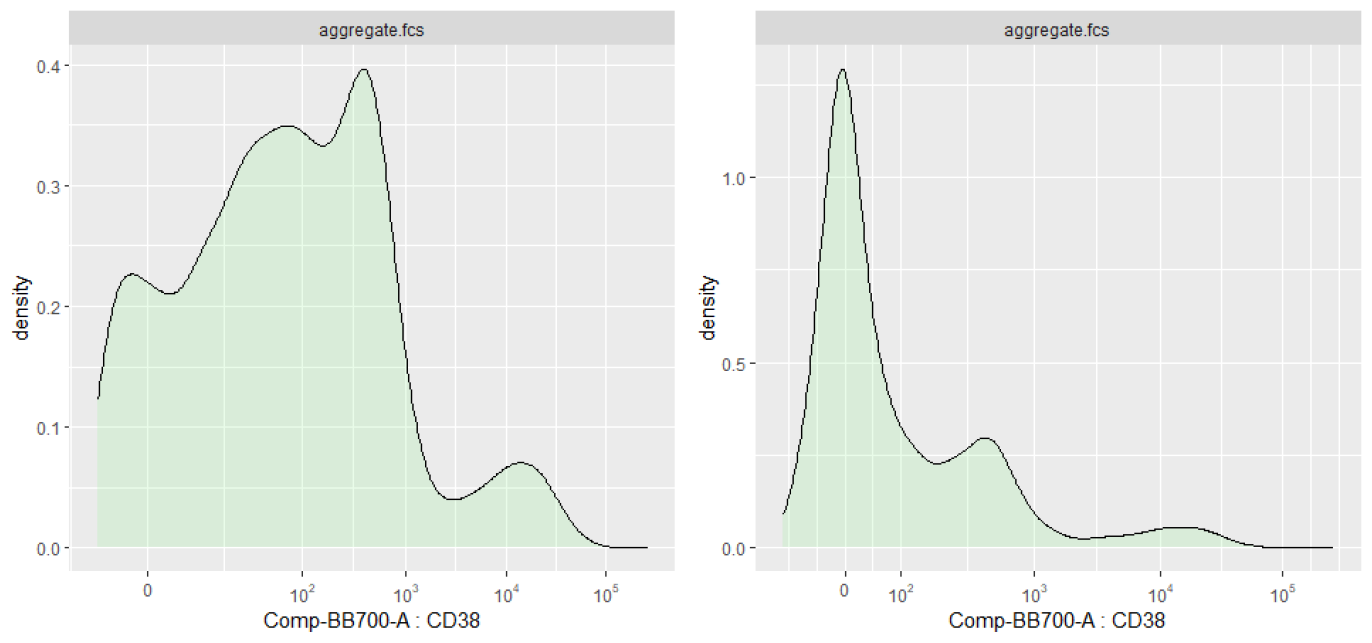
Manual parameters adjustment of the *logicle* transformation applied on marker CD38, for sample *D91_G01*. On the left, the density plot shows spurious oscillations in the negative domain. On the right, manual adjustment on the *positive decimals* parameter of the *logicle* transformation leads to a better looking transformed density, where one can more easily identify CD38-, CD38+ and CD38++ populations.

### Benchmarking results

As mentioned in the Methods section, we use *pipeComp* (Germain, Sonrel, and Robinson, 2020) to perform a benchmarking exercise, comparing two different pre-processing pipelines, i.e. the *PeacoQC*-based and the *flowAI*-based pipelines, on the 55 sample files of the *HBV chronic mouse* dataset, and calculating evaluation metrics in terms of how well the automated pipelines could match the manual pre-processing performed by an expert scientist (‘ground truth’). A global assessment shows comparable results between the two competing pipelines, consistently across all metrics (Figure 10). However, when directly contrasting sample by sample results (Figure 11) one can identify that the pipeline performance is rather heterogeneous across the 55 biological samples.

**Figure 10.**
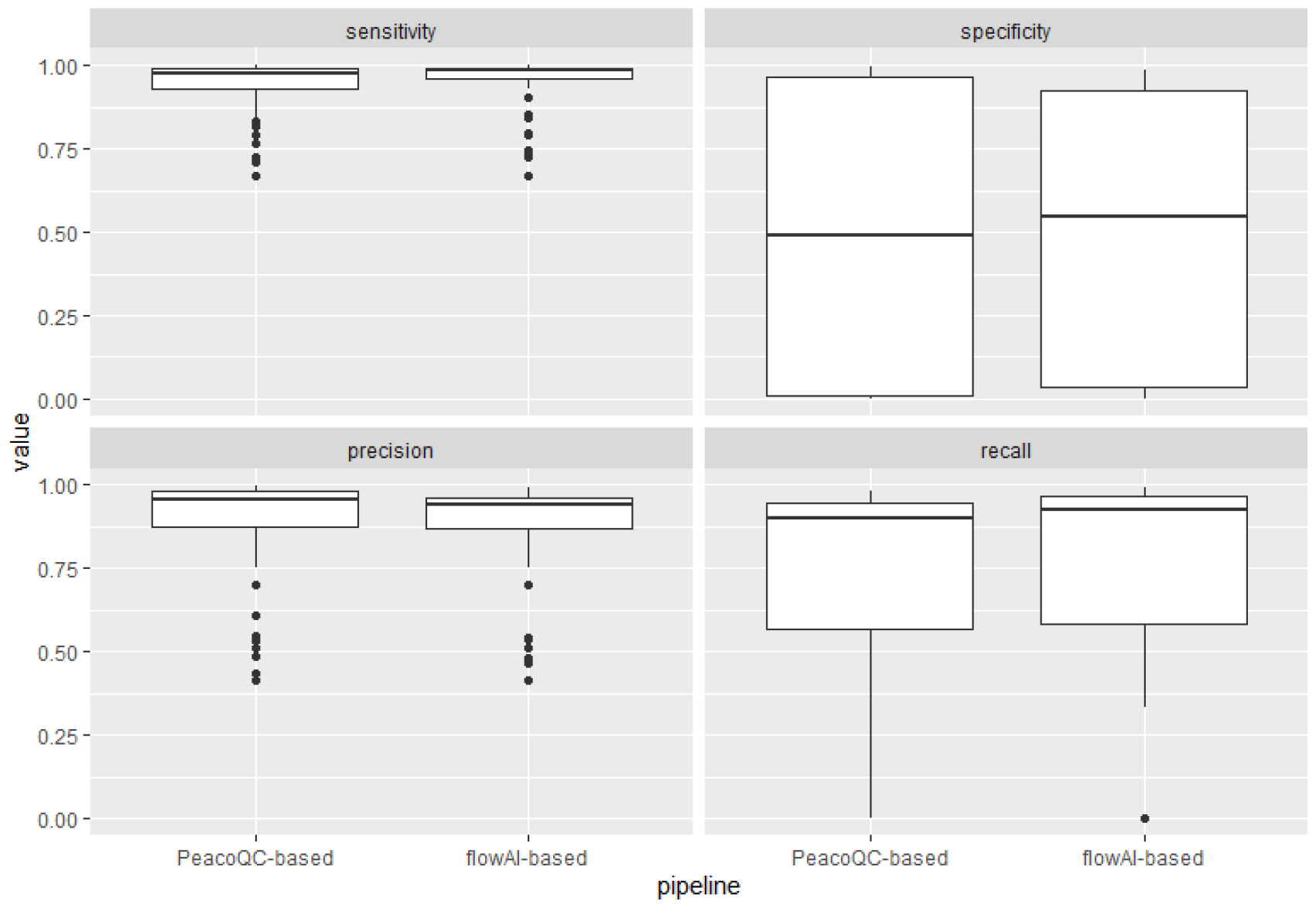
Box plots of the distributions of calculated evaluation metrics per sample, for the two competing pipelines. Globally, both pipelines perform very similarly, for all four evaluation metrics, i.e. sensitivity, specificity, precision and recall.

**Figure 11.**
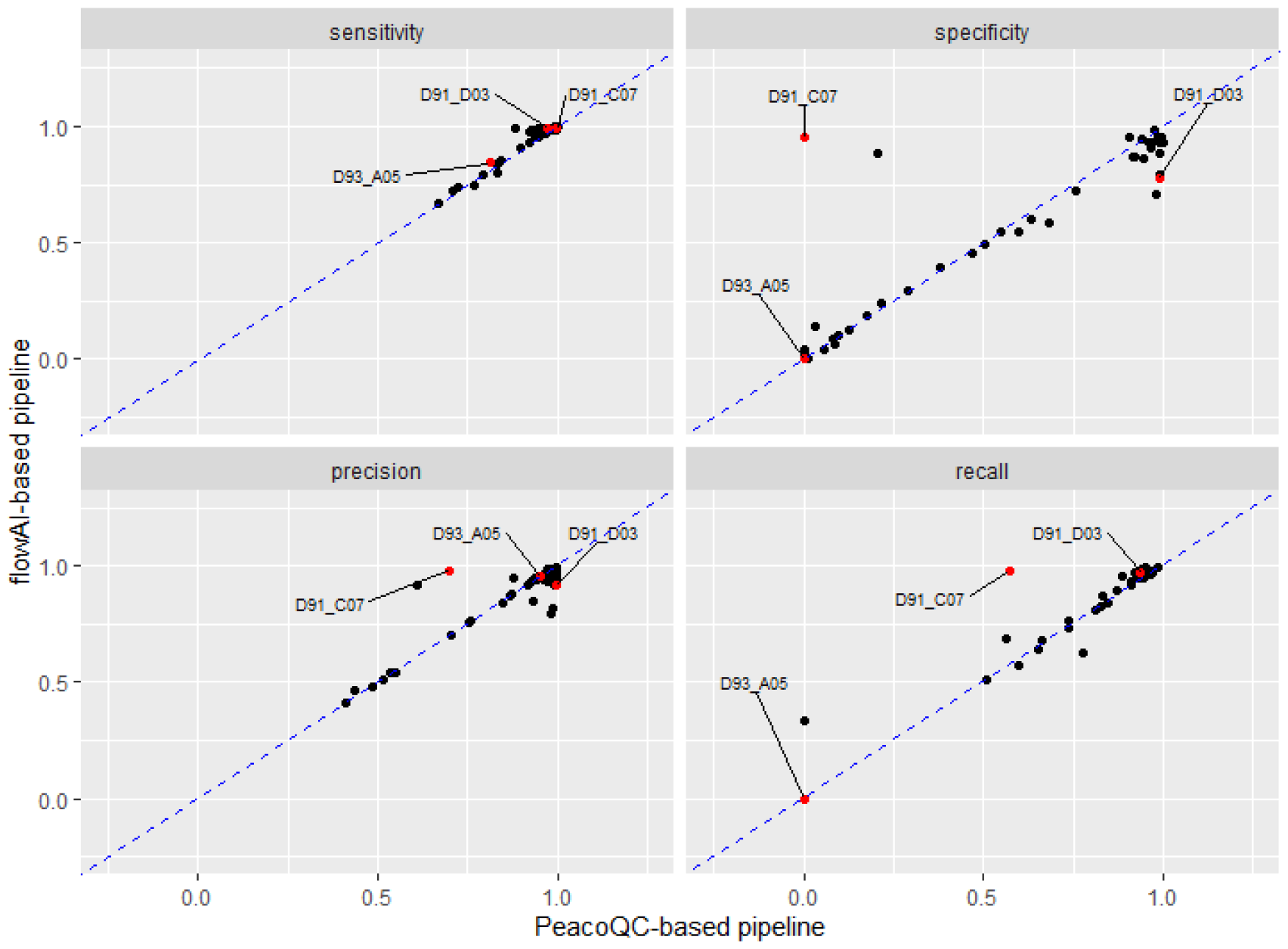
Scatter plots comparing the two pre-processing pipelines, each dot representing one of the 55 samples. Three specific samples are highlighted in red, corresponding to very different comparative behaviour of the two competing pipelines. Sample *D91_C07* is a unique sample for which the *flowAI*-based pipeline has a high specificity, but the *PeacoQC*-based pipeline has very low specificity. Sample *D93_A05* is one of the samples leading to low specificity for both pipelines, while *sample D91_D03* is representative of the samples for which both pipelines provide good specificity.

In order to better understand the behaviour of the two competing automatic pipelines on different samples, we selected three different samples, corresponding to different locations into the specificity plot of Figure 11. We then used *CytoPipelineGUI* to inspect the results at different steps, for the two automated pipelines as well as for the ‘ground truth’:

- Sample *D91_C07* was an outlier for which the *PeacoQC*-based pipeline obtained an almost zero specificity, while *flowAI*-based pipeline specificity was around an acceptable level of above 0.8. However, as shown in Figure 12, this was not due to the different *QC in time* algorithm (*PeacoQC* vs. *flowAI*), but to a lack of robustness of the dead cells removal algorithm, leading to an interaction phenomenon by which almost all events were removed in the dead cell removal step of the *PeacoQC*-based pipeline.
- Sample *D93_A05* resulted in a very low specificity for both pipelines. Investigation using *CytoPipelineGUI* revealed that this sample was in fact one of the low quality samples wherein the interesting cell population was a small minority of the events, while there was a great abundance of debris and dead cells (Figure S5). As a consequence, both pipelines were unable to automatically select the correct cell population, regardless of the *QC in time* method used.
- Sample *D91_D03* was an example where both automatic pipelines performed adequately without major issues. Here, the difference in metrics is effectively related to the choice of *QC in time* method. Looking at a specific visualization where time is displayed on the x axis (Figure S6), and based on both qualitative plot inspection and number of events comparison with the manual gating ground truth, *CytoPipelineGUI* reveals that *flowAI* method is too agressive in this case, while *PeacoQC* is too liberal.

**Figure 12.**
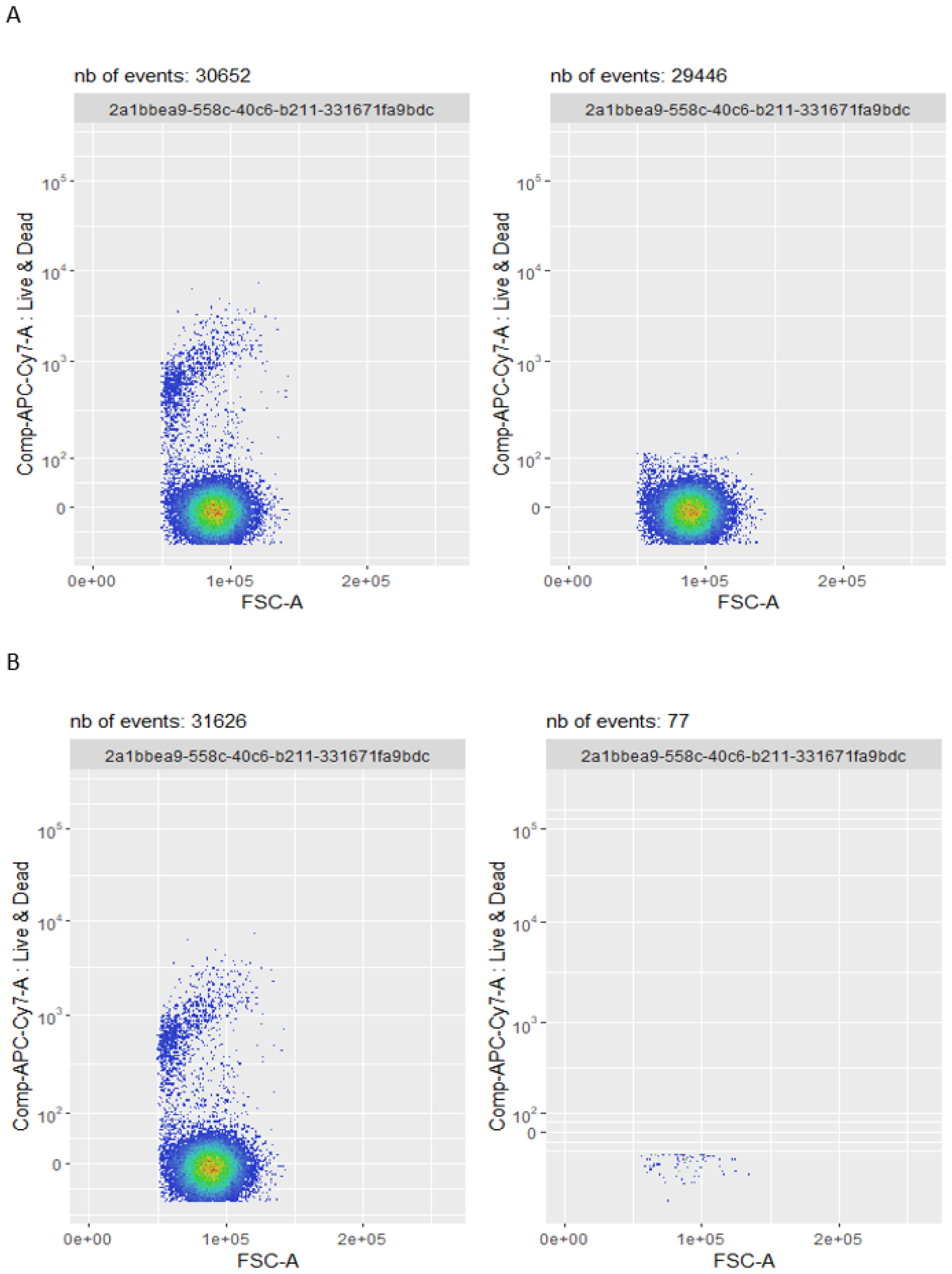
Comparison between the dead cells removal step between the *flowAI*-based pipeline (A), and the *PeacoQC*-based pipeline (B), on sample *D91_C07*. While the input set of events look very similar (left plots of panels A and B), the dead cells removal step of the *PeacoQC* pipeline (right plot of panel B) wrongly removes most of the events. This reveals a lack of robustness of the algorithm, unrelated to the*QC in time* method used (*flowAI* vs. *PeacoQC*).

Note that the conclusions of these visual inspections are particularly precious to the scientist in charge of building the data analysis pipelines, who is now able to get precise and accurate insight into why one pipeline performs better than the other, for specific samples. In particular, they are much better equipped to distinguish between an intrinsic performance difference between some competing methods, and surprising artefacts like a side effect of low sample quality or an interaction between two different steps.

## Discussion

### CytoPipeline, a flexible framework for building and running pre-processing pipelines

In this work, we have demonstrated the use of the *CytoPipeline* suite by implementing pre-processing pipelines on the *HBV chronic mouse* dataset. The implementation of *CytoPipeline*, with a centralized specification of the pipeline definition in a *json* file, leads to a better design of the pipeline code. As a result, we believe that the user productivity, when coding and testing different pipelines, can be greatly improved.

In order to illustrate this, we implemented the two *PeacoQC*-based and *flowAI*-based competing pipelines, described in Methods, in two *R* scripts, without using *CytoPipeline* objects, and looked into the duplication effort as well as the future extensibility of the code. These pieces of code are provided in the *2023-CytoPipeline-code* GitHub repository (see Code Availability in Declarations section).

Figure 13 provides a schematic comparison between these two pieces of code, as well as indicative number of code lines. Of course, these relates to one particular implementation, as there are countless ways to program the same pipelines. What is interesting to note, though, is that there is a high proportion of code duplication, but the differences are not only located in one single place, due to the subtle differences induced by the change of orders in the steps. This is likely to lead to a high code maintenance burden in the future, for instance when extending the program to many more pipeline instances, which can use different step methods, different method parameters etc. In constrast, let us recall that, when using *CytoPipeline*, the *R* code itself stays the same, as all differences are explicitly described in the input *json* file. This *json* file is easier to maintain and extend than the *R* scripts represented in Figure 13.

**Figure 13.**
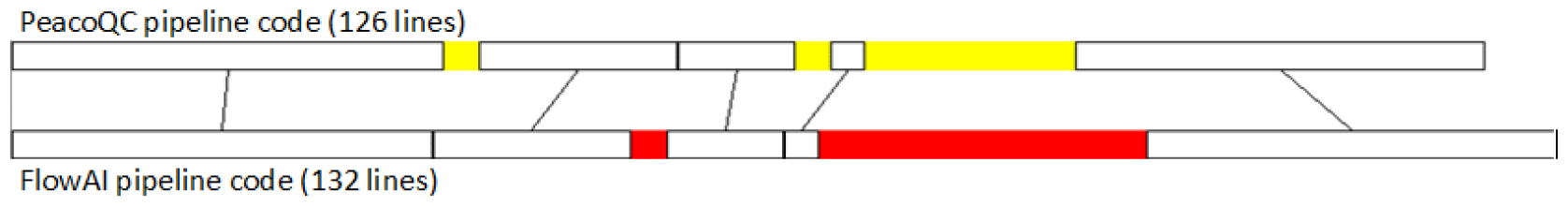
Structure of the *R* script implementations of the *PeacoQC*-based and *flowAI*-based pipelines. The common parts are shown in white, *PeacoQC*-based pipeline specific parts in yellow, and *flowAI*-based pipeline specific parts in red. Between the two pipelines, 79% of the code is in common, and the pipeline specific parts are not fully gathered in one single location.

### CytoPipeline provides a standardized and user-friendly tool for visual investigations

We have presented a series of use cases of *CytoPipeline* visualizations. In all these use cases, we took advantage of the same set of visualization tools, in a standardized way, but translated into different contexts, whatever the underlying methods used for the pre-processing pipelines. Also during the investigation of the benchmarking results, visual comparisons could be made with a ground truth manual gating, again using the same tools. Besides, the interactive GUI applications, implemented in *CytoPipelineGUI*, provide user interactivity and facilitate the investigation process.

As stated in the introduction, these visual assessments are extremely important for the scientists, as they provide a unique mean to:

- visually control for the quality of the data samples, and acquire insight on the corresponding sample variability;
- visually check the robustness of the methods used in a given pre-processing pipeline, including the adequacy of the chosen user input parameters;
- visually compare different pre-processing pipeline settings. This can range from comparing different possible choices of method for a particular step, to assessing which one of two or more competing pipelines, possibly mixing different step methods in different orders, is performing better for the considered dataset.

### CytoPipeline allows user intuitive insight into benchmarking results

As part of this work, we have implemented a benchmarking comparing two competing pre-processing pipelines, with the main objective of showing the benefits of using *CytoPipeline* visualization tools, as a complement to the benchmarking itself. We showed that detailed comparison plots help the user investigating some specific benchmarking results, hence getting better intuition into the benchmarking outcome. We have indeed demonstrated that there can be numerous reasons why a pipeline instance performs better than another on specific samples, and it is key for the scientist to get a clear view of these reasons, and their possible links with sample characteristics. Therefore, we think that *CytoPipeline* is a powerful tool for interpreting the outcome of benchmarking studies.

### Using the proportion of events kept at each step as a diagnostic tool

As was shown in various figures in the Results section (see e.g. Figure 12), *CytoPipelineGUI* computes the number of events that are retained at each step (shown as subtitles in the individual density plots). Tracking these changes throughout the pre-processing steps of a pipeline for different samples is a useful quality control. This can be implemented using some of the *CytoPipeline* functions, and is shown on Figure 14.

**Figure 14.**
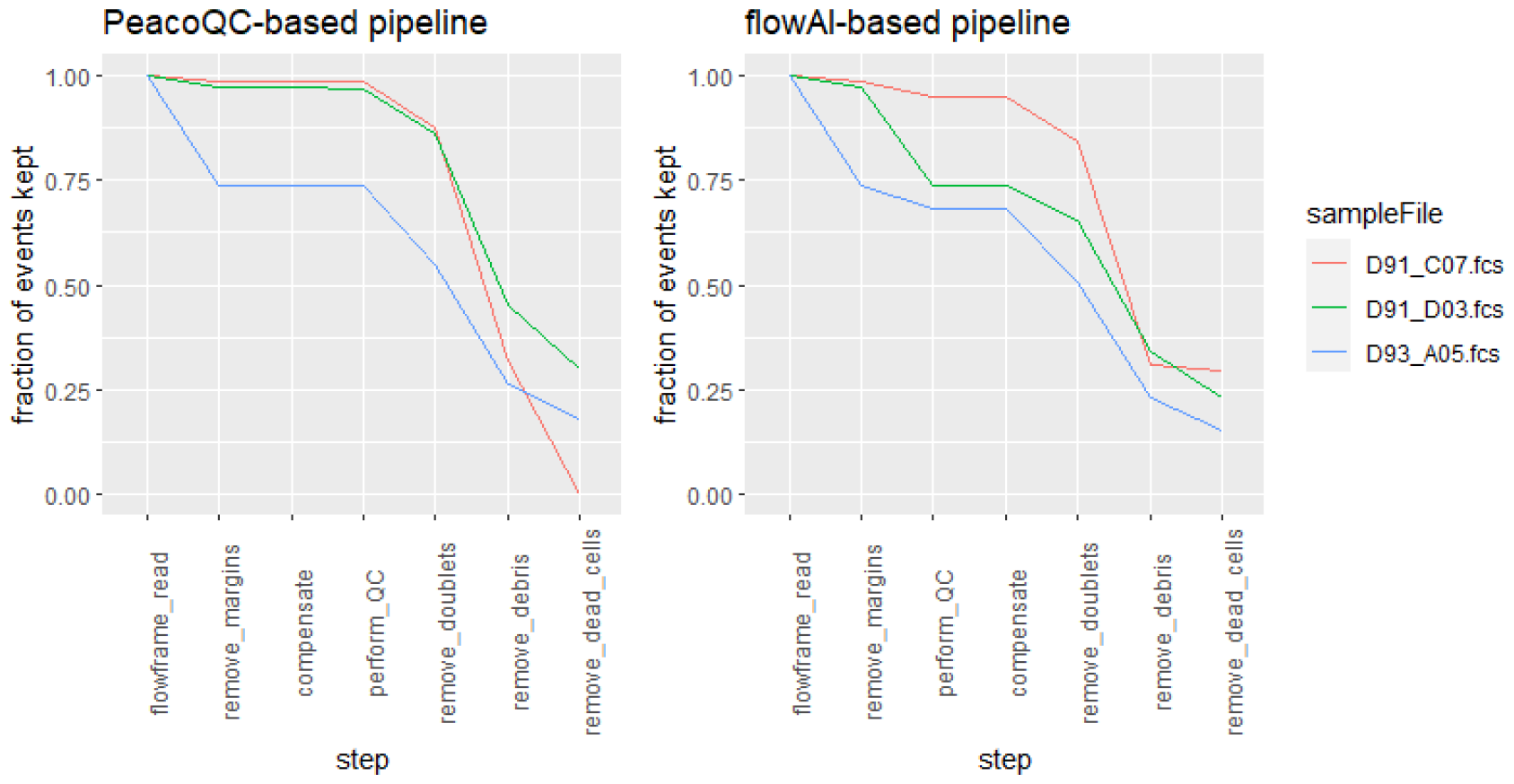
Plots showing the proportion of retained events at each pre-processing step, for each sample. On the left, the *PeacoQC*-based pipeline shows, for sample *D91_C07*, a sharp drop in the last *remove_dead_cells* step. On the right, the *flowAI*-based pipeline does not show the same phenomenon.

### Limitations and possible extensions of the work

The *CytoPipeline* suite of *R* packages can be positioned as a tool to facilitate the design, testing and comparison of pre-processing pipelines for the end user. It is not meant to be:

- A novel pre-processing pipeline in itself, as it does not provide new methods for the various pre-processing steps (although it includes some functions calling some widely used methods), nor an innovative way to combine some of these.
- A tool facilitating benchmarking automation, like *pipeComp*. For example, unlike *pipeComp* (Germain, Sonrel, and Robinson, 2020), *CytoPipeline* does not provide any optimization solution to reduce the amount of CPU time and memory to run a potentially huge amount of (combinations of) possible pipelines. However, as mentioned before, *CytoPipeline* is used to facilitate the interpretation of results produced with benchmarking tools.

Regarding scalability, one should distinguish CPU and memory from hard drive storage requirements. CPU- and memory-wise, *CytoPipeline* has no particular issues when dealing with large number of samples, as long as each single fcs file can fully reside in memory. Indeed, as described in the Methods section, the engine that executes pre-processing pipelines supports both sequential and parallel file processing, and benefits from all multi-tasking scheduling options provided by the *BiocParallel* Morgan et al., 2023 package. However, storage-wise, caching data at each step leads to large storage needs when processing many files. Typically, when analysing datasets including hundreds of fcs files, with several millions of events, compared across several pipelines and many processing steps, storage needs can require several terabytes. In those cases, users of *CytoPipeline* will typically need to call on high capacity storage facilities.

Another limitation of our work is the following: while *CytoPipelineGUI* is a powerful visualization tool for exploring specific pipeline steps for one or two samples, it does not provide an overall quality control of all samples at once. In that sense, it would be of useful, especially for large datasets, to provide a global view of how samples differ at each pre-processing step. As mentioned above, one such diagnostic view can be obtained, by plotting the fraction of retained events at each preprocessing step (Figure 14). Another promising approach focuses on the visualisation of all samples at once to identify specific outliers Hauchamps and Gatto, 2024.

Finally, another possible extension would be to further develop *CytoPipeline*, as to not only include the building and assessment of pre-processing steps, but also include support for subsequent steps of the data analysis: batch correction, population identification, etc.

## Conclusion

In this work, we have introduced a suite of *R* packages, *CytoPipeline* and *CytoPipelineGUI*, that helps building, visualizing and assessing pre-processing pipelines for flow cytometry data. We have demonstrated several use cases on a real life dataset, and highlighted several concrete benefits of these tools. For the new user, the packages come with ample documentation and tutorial videos, accessible through the package vignettes. We trust that using *CytoPipeline* will favour productivity in testing and assessing alternative data pre-processing pipelines, with the aim of designing good pre-processing and QC solutions for each particular context. The latter can be the specific type of biological sample, technology used (conventional flow cytometry, cytof, spectral flow cytometry), panel composition, experimental design etc., which in turn highly depend on the biological question at hand.

## Declarations

### Funding

This work was funded by GlaxoSmithKline Biologicals S.A., under a cooperative research and development agreement between GlaxoSmithKline Biologicals S.A. and de Duve Institute (UCLouvain).

## Competing Interests

- B.B., S.D., M.H., M.T., S.T., and D.L. are employees of the GSK group of Companies.
- B.B., S.D., M.T., S.T., and D.L. report ownership of GSK shares.
- B.B., M.T., and S.T. are listed as inventors on patent(s) owned by the GSK group of companies.
- P.H. is a student at the de Duve Institute (UCLouvain) and participates in a post graduate studentship program at GSK.
- L.G. reports no competing interest.

## Author contributions

- Conceptualization: P.H., L.G.;
- Methodology: P.H., L.G., D.L., S.D.;
- Software: P.H., L.G.;
- Data collection: B.B., M.H.;
- Writing - original draft: P.H., B.B.;
- Writing - review & editing: L.G., D.L., S.T., S.D., M.H., M.T.;
- Supervision: L.G., D.L., S.T., M.T.

## Availability of data and materials

Raw flow cytometry data files, as well as the manual gating information considered as the ground truth for the benchmarking, are available on Zenodo (DOI:10.5281/zenodo.8425840).

## Code availability

All code needed to reproduce the results presented in the current article is available on the following GitHub repository: https://github.com/UCLouvain-CBIO/2023-CytoPipeline-code, of which a release has been archived on Zenodo (DOI:10.5281/zenodo.8425840).

## Acknowledgment

This preprint was created using the LaPreprint template (https://github.com/roaldarbol/lapreprint) by Mikkel Roald-Arbøl.

## Supplementary Materials

### Supplementary Tables

**Table S1.** Scale transformation steps, common to both *PeacoQC*-based and *flowAI*-based pipelines.

| Step | Name | R package | Version | Method |
| --- | --- | --- | --- | --- |
| 1 | flowframe_read | *flowCore* | 2.12.0 | *read.flowSet()* |
| 2 | remove_margins | *PeacoQC* | 1.10.0 | *removeMargins()* |
| 3 | compensate | *flowCore* | 2.12.0 | *compensate()* |
| 4 | flowframe_aggregate | *CytoPipeline* | 1.0.0 | *aggregateAndSample()* |
| 5 | scale_transform_estimate | *flowCore* | 2.12.0 | *estimateLogicle()* |

**Table S2.** *PeacoQC*-based pipeline pre-processing steps.

| Step | Name | R package | Version | Method |
| --- | --- | --- | --- | --- |
| 1 | flowframe_read | *flowCore* | 2.12.0 | *read.FCS()* |
| 2 | remove_margins | *PeacoQC* | 1.10.0 | *removeMargins()* |
| 3 | compensate | *flowCore* | 2.12.0 | *compensate()* |
| 4 | perform_QC | *PeacoQC* | 1.10.0 | *PeacoQC()* |
| 5 | remove_doublets | *CytoPipeline* | 1.0.0 | *removeDoubletsCytoPipeline()* |
| 6 | remove_debris | *flowClust* | 3.38.0 | *tmixFilter()* |
| 7 | remove_dead_cells | *flowDensity* | 1.34.0 | *deGate()* |

**Table S3.** *flowAI*-based pipeline pre-processing steps.

| Step | Name | R package | Version | Method |
| --- | --- | --- | --- | --- |
| 1 | flowframe_read | *flowCore* | 2.12.0 | *read.FCS()* |
| 2 | remove_margins | *PeacoQC* | 1.10.0 | *removeMargins()* |
| 3 | perform_QC | *flowAI* | 1.30.0 | *flow_auto_qc()* |
| 4 | compensate | *flowCore* | 2.12.0 | *compensate()* |
| 5 | remove_doublets | *CytoPipeline* | 1.0.0 | *removeDoubletsCytoPipeline()* |
| 6 | remove_debris | *flowClust* | 3.38.0 | *tmixFilter()* |
| 7 | remove_dead_cells | *flowDensity* | 1.34.0 | *deGate()* |

### Supplementary Figures

**Figure S1.**
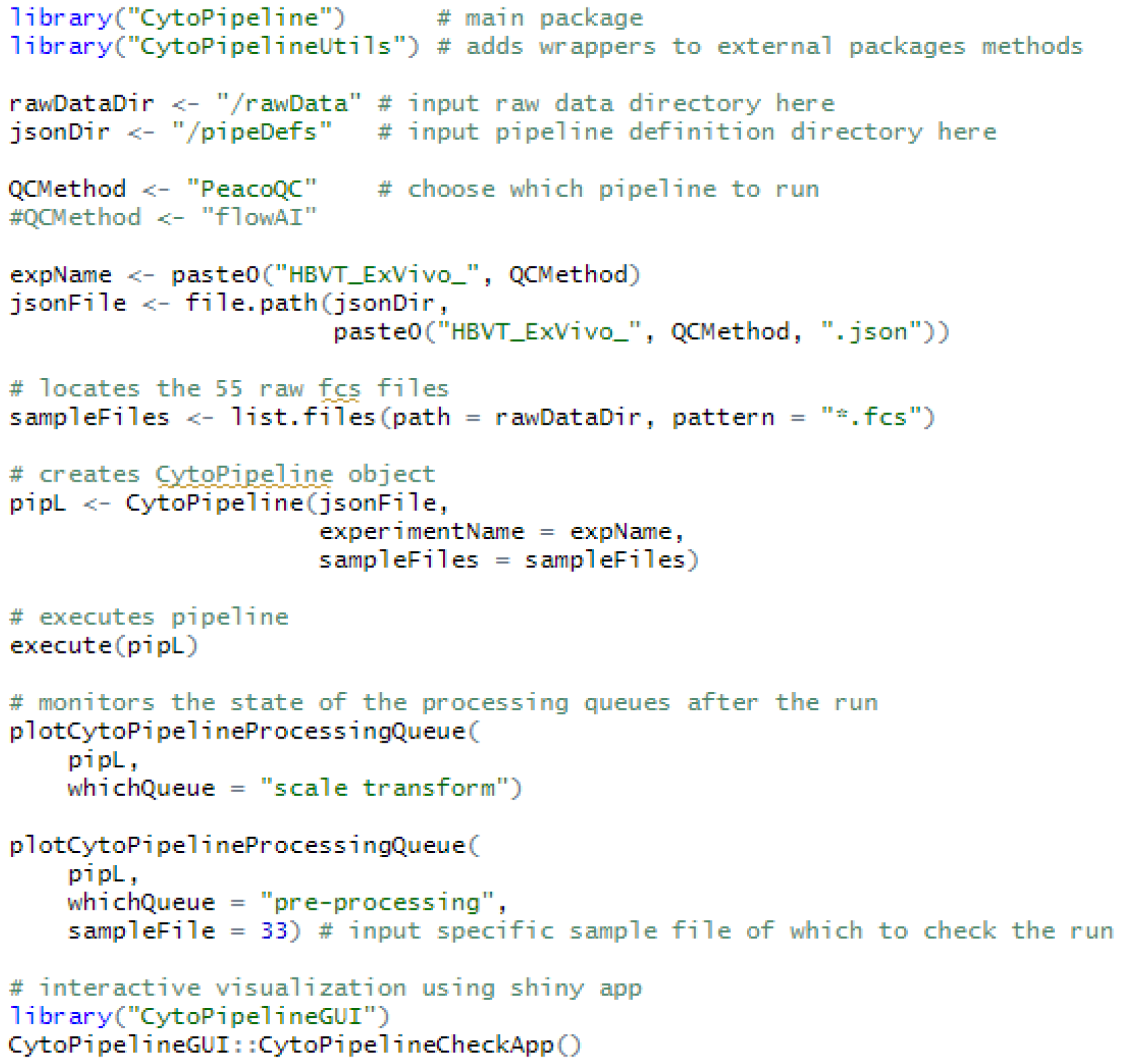
Simple sample *R* code to define a *CytoPipeline* object, run it and visualize the results.

**Figure S2.**
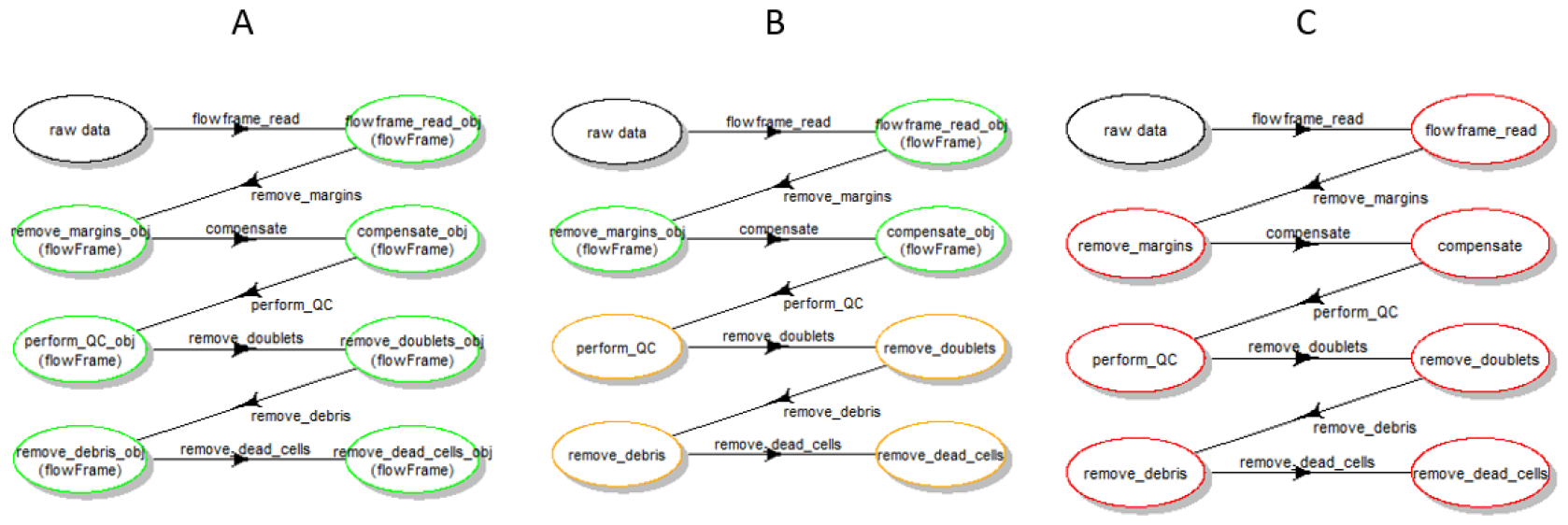
Illustration of the colour code used to represent the pipeline workflows. Green steps are the ones that have already been run and produced results, indicating a non problematic run. Orange steps are the ones that are correctly defined, but have not run yet for the displayed sample file. Red steps show an inconsitency problem between the already stored results, and the *CytoPipeline* object, meaning that the pipeline was previously run, either with a different number of steps, or with one or several steps that used a different method or different set of parameters. On the above picture, part *A* shows a situation where part of the pipeline has fully run. Part *B* shows a situation of pipeline that has run, but not all steps. Part *C* shows an inconsistency between the pipeline definition and the results stored.

**Figure S3.**
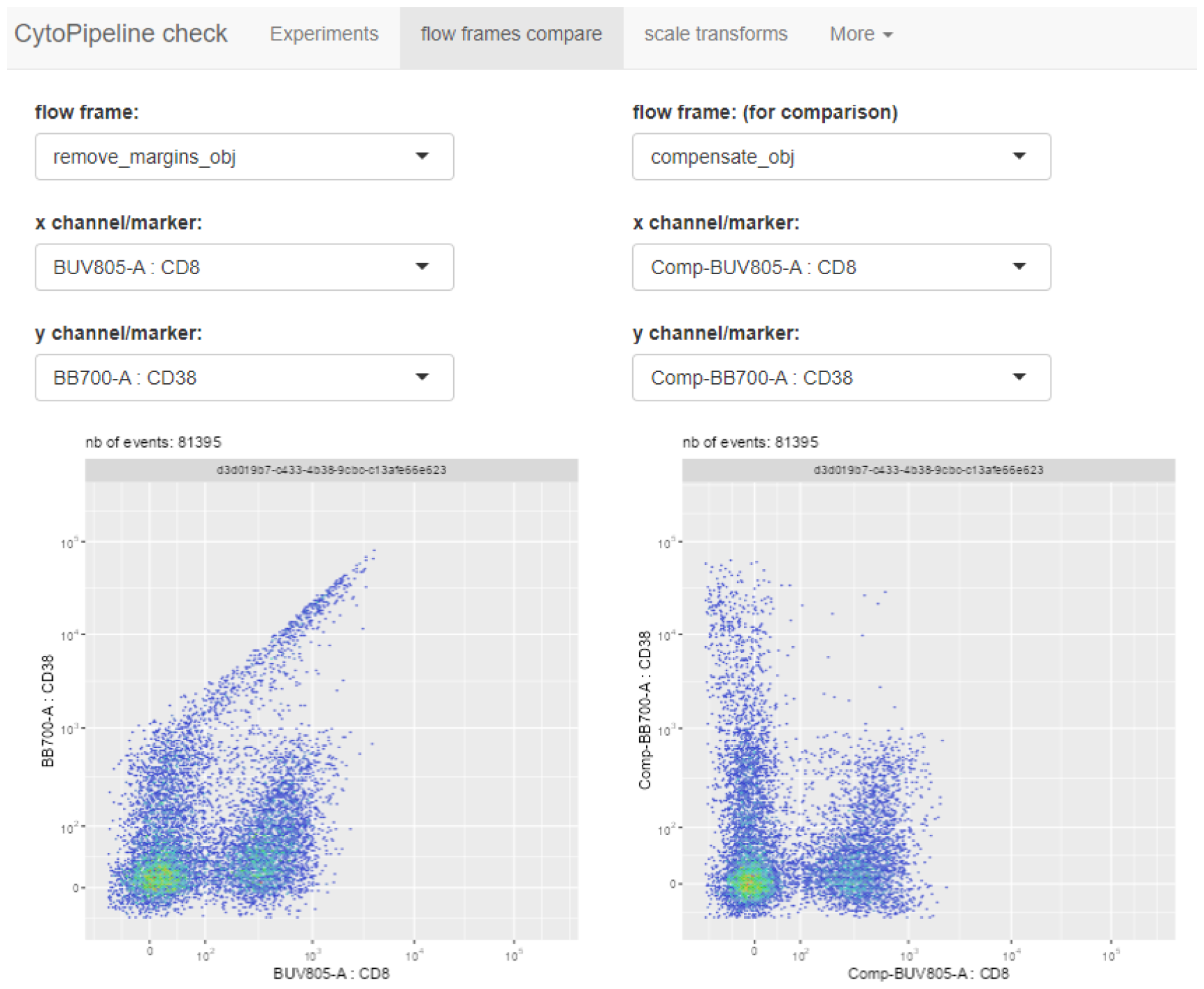
Screenshot of the interactive GUI application allowing to compare flow frames, and implemented in the *CytoPipelineGUI* package. Here the displayed plots correspond to Figure 4 of the main text.

**Figure S4.**
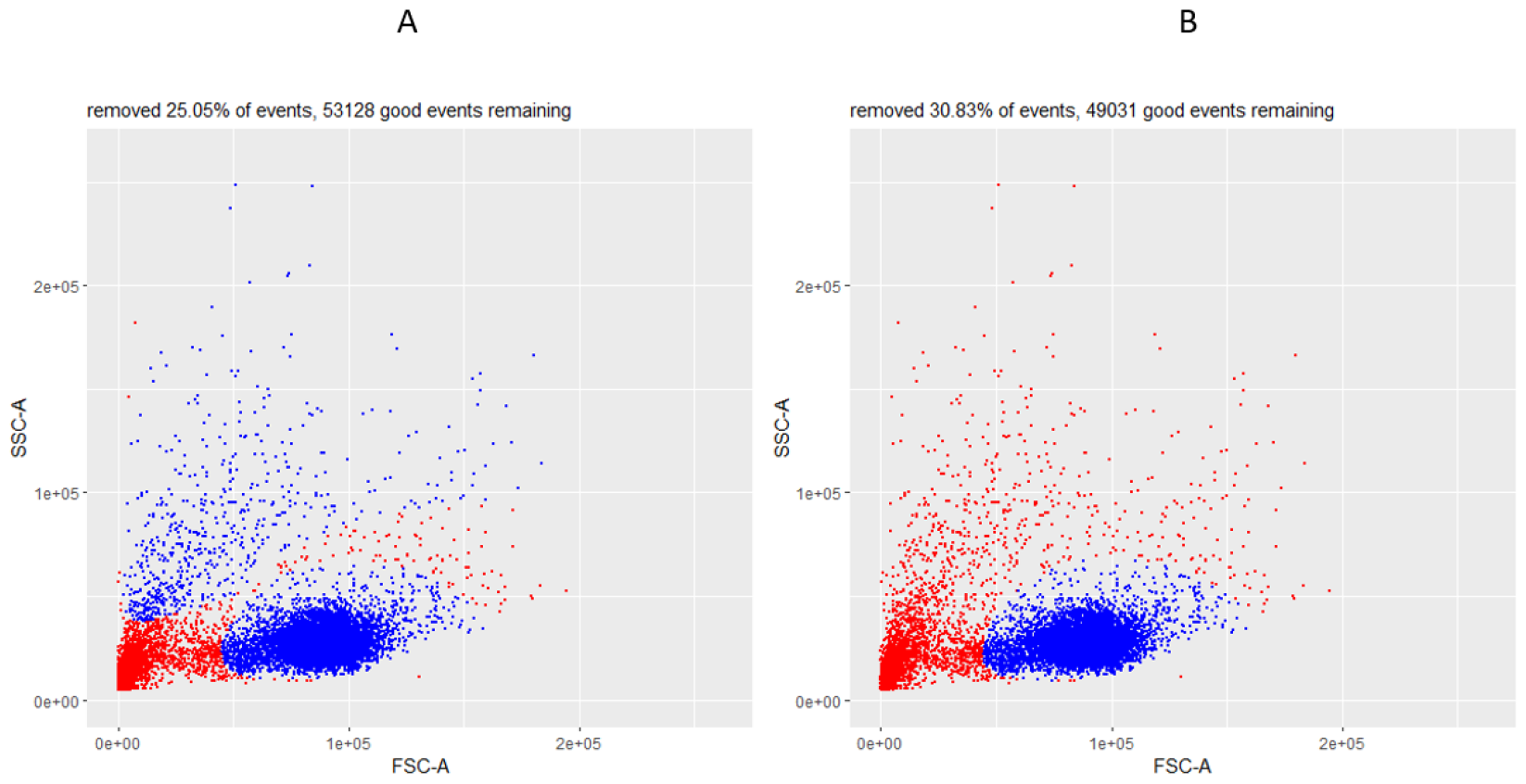
Comparison between the outcome of the debris removal step between the *PeacoQC*-based pipeline with 3 clusters (A), and the *PeacoQC*-based pipeline with 2 clusters (B). The latter setting better eliminates undesirable events than the former.

**Figure S5.**
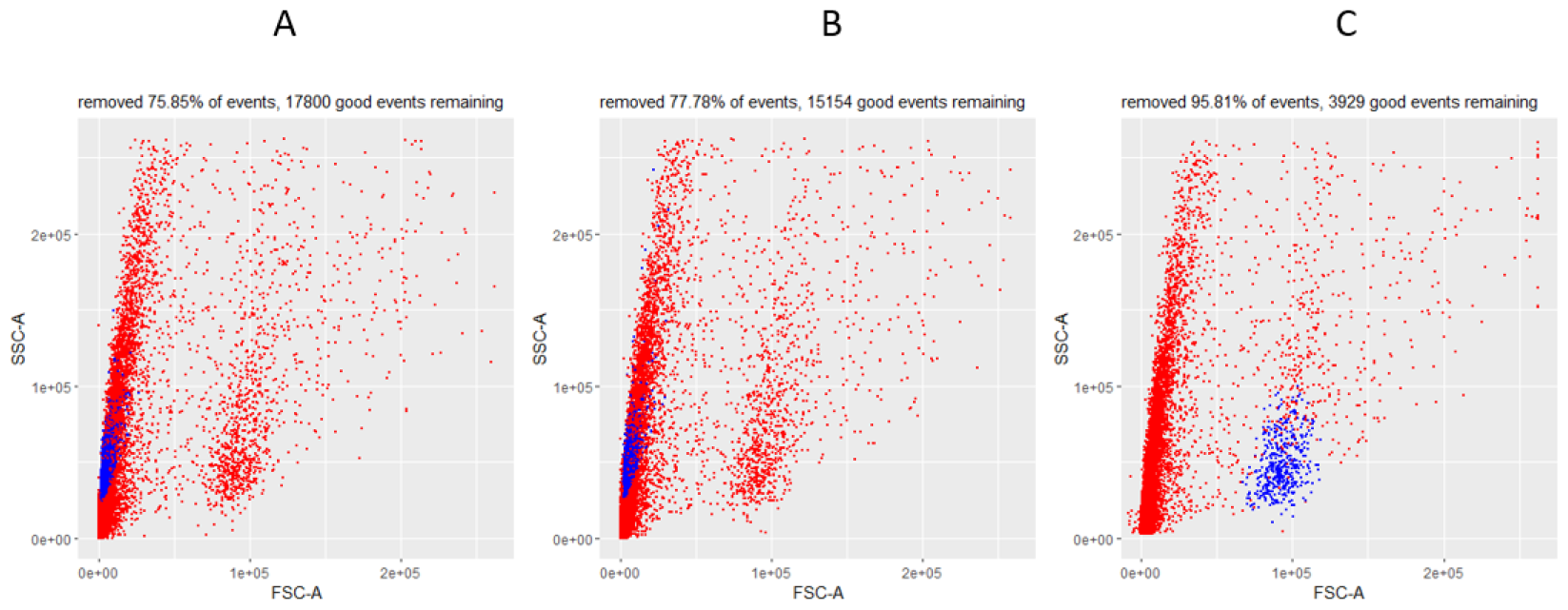
Comparison between the outcome of three combined steps (doublets, debris and dead cells removals) between the *PeacoQC*-based pipeline (A), the *flowAI*-based pipeline (B), and the ground truth (C), on sample *D93_A05*. Due to the bad quality of this sample, i.e. the majority of the events were composed of debris and dead cells, both automatic pipelines were unable to select the right population of lymphocytes.

**Figure S6.**
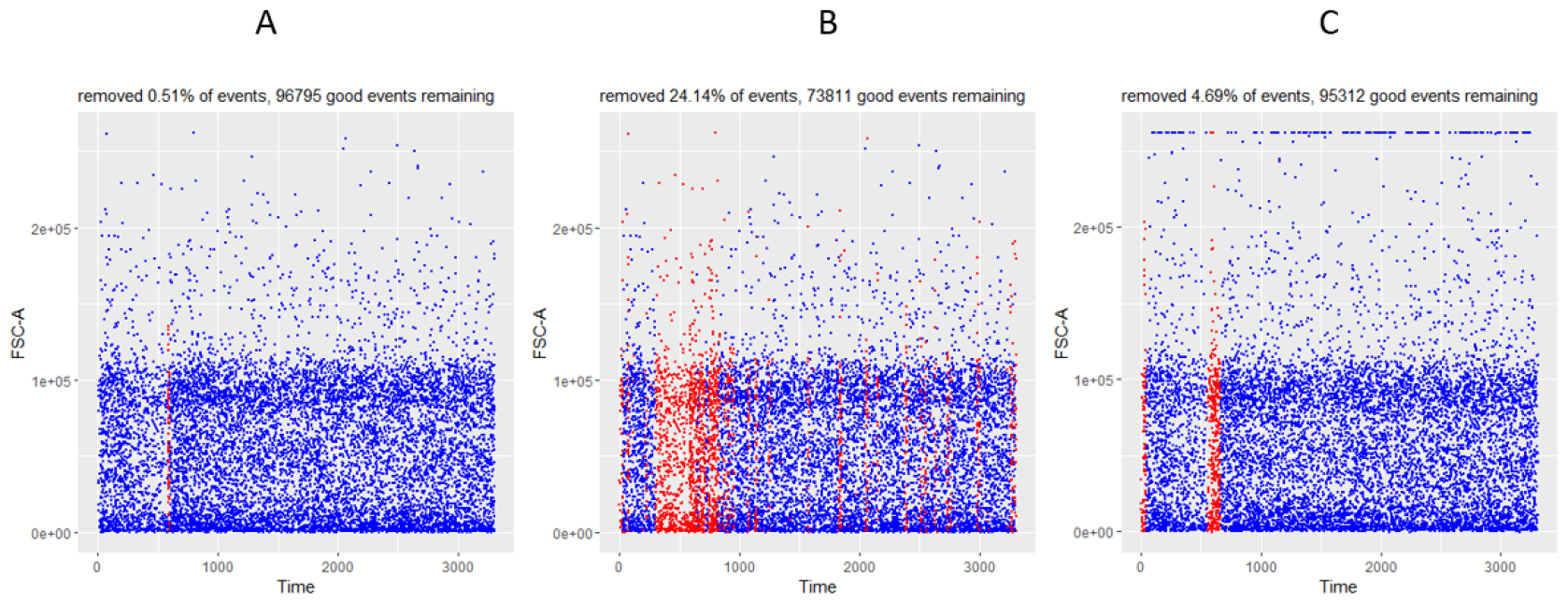
Comparison between the *QC in time* step with *PeacoQC* (A), with *flowAI* (B), and the time gate applied in manual gating (ground truth) (C), on sample *D91_D03. flowAI* removes more event than the ground truth, while *PeacoQC* removes less events.

### Supplementary text file 1: json file describing the *PeacoQC*-based pre-processing pipeline

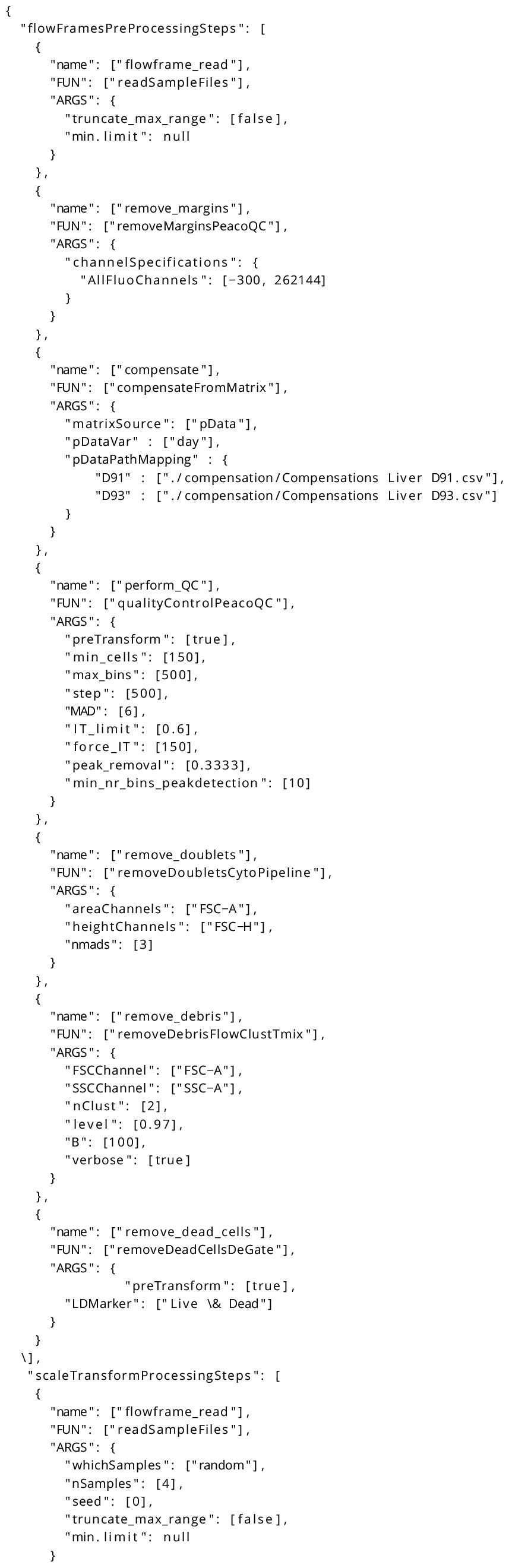
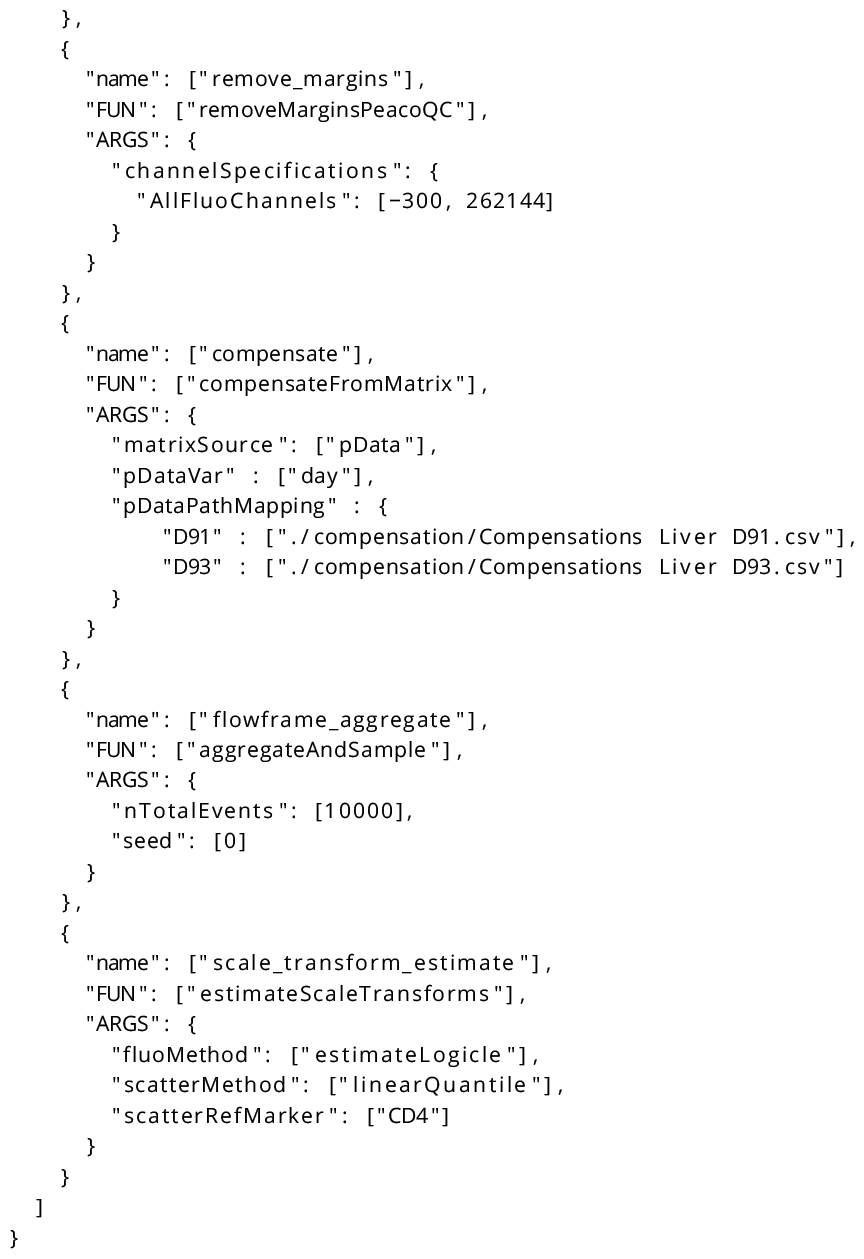

### Supplementary text file 2: json file describing the *flowAI*-based pre-processing pipeline

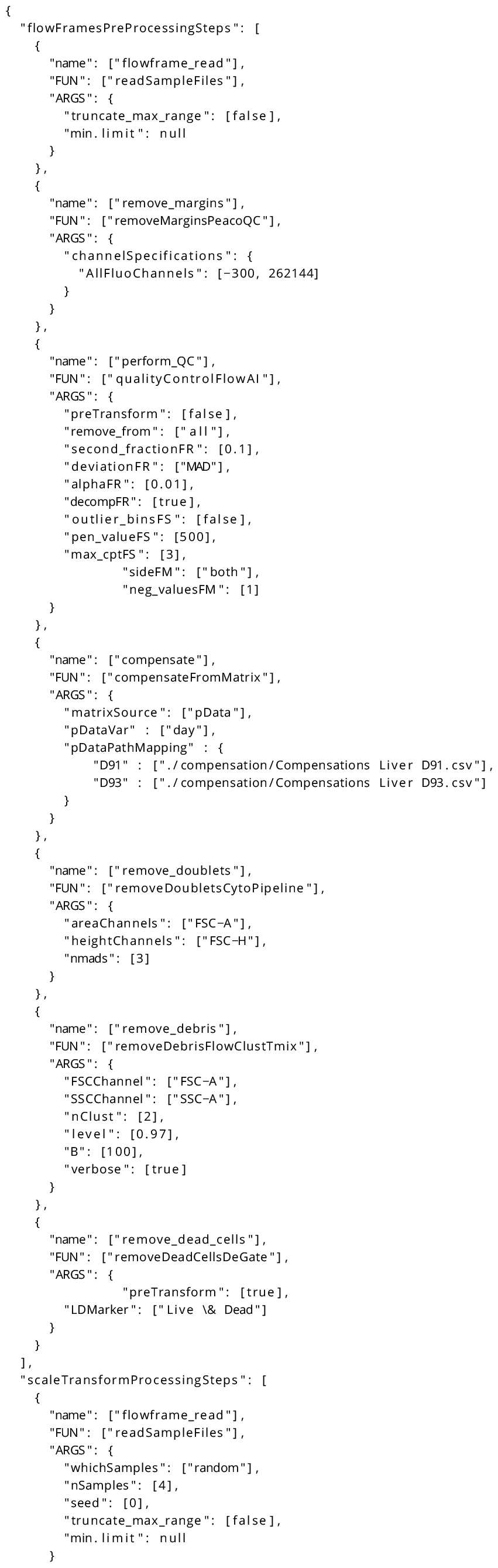
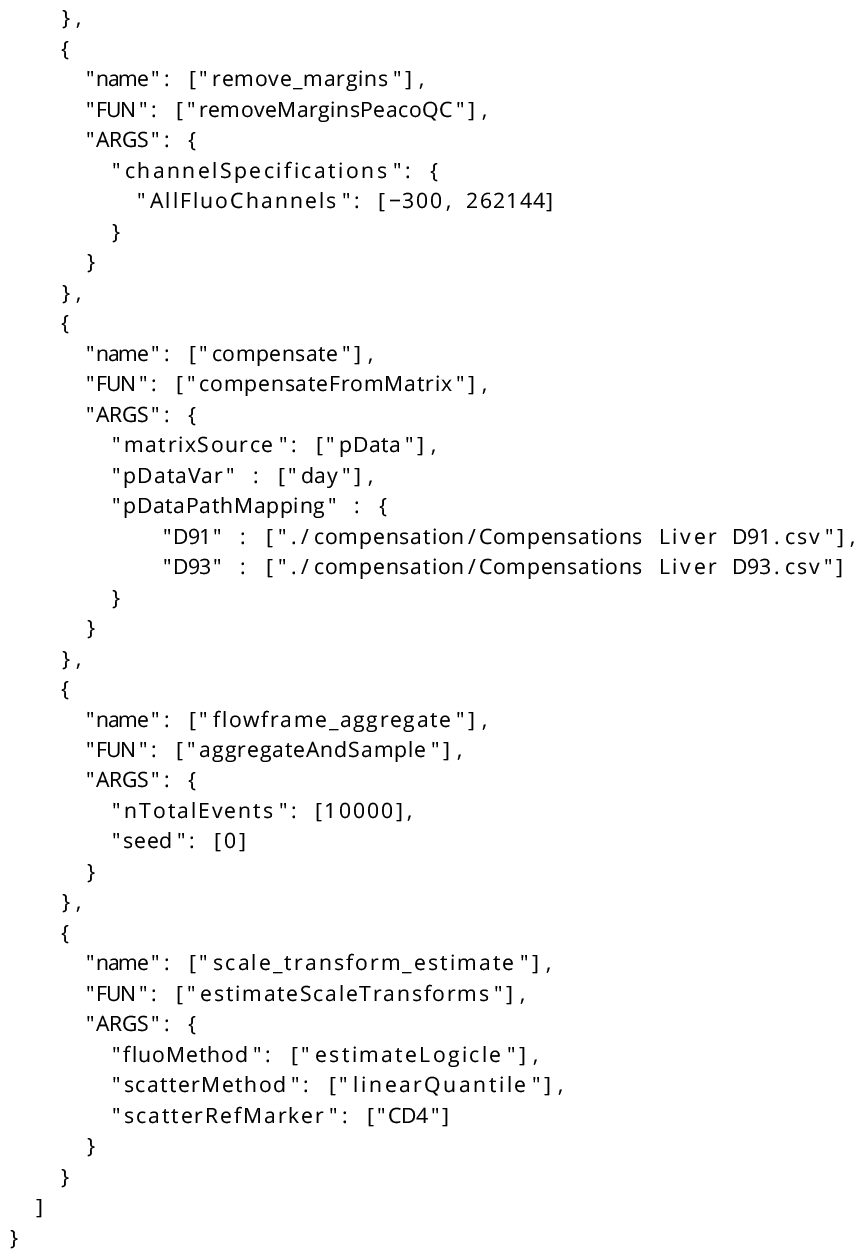

## Footnotes

1 note that *CytoPipeline* also provides methods to define a pipeline and its steps programmatically in *R*, without providing a text file as an input.

## Notes

https://github.com/UCLouvain-CBIO/2023-CytoPipeline-code

### Summary of Updates

- Added proportion of events across pre-processing steps. - Discussion on scalability. - Other minor updates.

